# Complexity of enhancer networks predicts cell identity and disease genes revealed by single-cell multi-omics analysis

**DOI:** 10.1101/2022.05.20.492770

**Authors:** Danni Hong, Hongli Lin, Lifang Liu, Muya Shu, Jianwu Dai, Falong Lu, Mengsha Tong, Jialiang Huang

**Author notes:** To whom correspondence should be addressed. E-mail: Jialiang Huang. These authors contributed equally (Danni Hong, Hongli Lin).

## Abstract

Many enhancers exist as clusters in the genome and control cell identity and disease genes; however, the underlying mechanism remains largely unknown. Here, we introduce an algorithm, eNet, to build enhancer networks by integrating single-cell chromatin accessibility and gene expression profiles. Enhancer network is a gene regulation model we proposed that not only delineates the mapping between enhancers and target genes, but also quantifies the underlying regulatory relationships among enhancers. The complexity of enhancer networks is assessed by two metrics: the number of enhancers and the frequency of predicted enhancer interactions (PEIs) based on chromatin co-accessibility. We apply eNet algorithm to a human blood dataset and find cell identity and disease genes tend to be regulated by complex enhancer networks. The network hub enhancers (enhancers with frequent PEIs) are the most functionally important in enhancer networks. Compared with super-enhancers, enhancer networks show better performance in predicting cell identity and disease genes. The establishment of enhancer networks drives gene expression during lineage commitment. Applying eNet in various datasets in human or mouse tissues across different single-cell platforms, we demonstrate eNet is robust and widely applicable. Thus, we propose a model of enhancer networks containing three modes: Simple, Multiple and Complex, which are distinguished by their complexity in regulating gene expression.

Taken together, our work provides an unsupervised approach to simultaneously identify key cell identity and disease genes and explore the underlying regulatory relationships among enhancers in single cells, without requiring the cell type identity in advance.

**Highlights:**

- eNet, a computational method to build enhancer network based on scATAC-seq and scRNA-seq data
- Cell identity and disease genes tend to be regulated by complex enhancer networks, where network hub enhancers are functionally important
- Enhancer network outperforms the existing models in predicting cell identity and disease genes, such as super-enhancer and enhancer cluster
- We propose a model of enhancer networks in gene regulation containing three modes: Simple, Multiple and Complex

## Introduction

Enhancers play a central role in orchestrating spatiotemporal gene expression programs during development and diseases (Consortium, 2012; Long et al., 2016; Maurano et al., 2012). Many enhancers exist as clusters in the genome to control gene expression, termed enhancer clusters, which control the expression of the same target gene (Blobel et al., 2021). Enhancer clusters are remarkably widespread features in the genome and provide an effective regulatory buffer for phenotypic robustness during development (Osterwalder et al., 2018; Perry et al., 2011). Several enhancer clusters in the genome have been described as super-enhancers (SEs), which exhibit disproportionately high signals for enhancer marks and control the expression of genes that define cell identity and diseases (Hnisz et al., 2013).

Genome editing using the CRISPR/Cas9 system offers an opportunity for dissecting the functions of enhancer clusters (Jinek et al., 2012). Several groups, including ours, have utilized genome editing assays to functionally dissect individual constituent elements of a couple of SEs (Bahr et al., 2018; Cai et al., 2020; Canver et al., 2015; Fulco et al., 2016; Hay et al., 2016; Huang et al., 2016; Kai et al., 2021; Shin et al., 2016; Thomas et al., 2021). These studies suggest the diversity of enhancer cluster regulatory mechanisms, where the individual components may act additively, redundantly, synergistically, or temporally. Meanwhile, genome-wide chromatin conformation information has been used to investigate the relationship among the individual components of enhancer clusters and their effects on target gene expression (Dixon et al., 2012; Lieberman-Aiden et al., 2009; Liu et al., 2017; Rao et al., 2014; Schoenfelder and Fraser, 2019; Song et al., 2020). We and other groups uncover hub enhancers, the enhancers with frequent chromatin interactions, play distinct roles in chromatin organization and gene activation (Huang et al., 2018; Huang et al., 2015; Liu et al., 2020; Schmitt et al., 2016).

Single-cell sequencing techniques have been developed to measure molecular heterogeneity among individual cells, such as single cell RNA sequencing (scRNA-seq) for gene expression and single-cell Assay for Transposase-Accessible Chromatin using sequencing (scATAC-seq) for chromatin accessibility (Buenrostro et al., 2015; Tang et al., 2009). It can even be used to achieve simultaneous detection of chromatin accessibility and gene expression in the same cells (Cao et al., 2018; Chen et al., 2019; Ma et al., 2020; Zhu et al., 2019). A large amount of single cell multi-omics profiles of chromatin accessibility and gene expression have been generated in various biological systems (Argelaguet et al., 2019; Granja et al., 2019; Li et al., 2021; Sarropoulos et al., 2021; Trevino et al., 2021; Ziffra et al., 2021), thereby providing ample opportunities to further understand the functions and mechanisms of enhancer clusters in single cells. For example, the co-accessible pairs of DNA elements predicted by Cicero from scATAC-seq data correspond with the chromatin contacts captured via ChIA-PET or promoter capture Hi-C (Pliner et al., 2018). However, these existing studies have largely focused on connecting enhancers with their target genes, but rarely on the regulatory relationship between enhancers. There remains a lack of method development to quantitatively assess how individual elements work together to regulate gene expression.

In this study, we proposed the concept of enhancer network complexity, which not only connects enhancers to putative target genes, but also quantifies how multiple enhancers interact with each other to regulate precise gene expression. Briefly, we developed a computational method termed eNet to build enhancer networks based on single-cell chromatin accessibility and gene expression data. Applying eNet on various biological systems, we found that the complexity of enhancer networks can predict cell identity and disease genes. Overall, we proposed a model of enhancer networks, which is not only useful in predicting cell identity and disease genes, but also provides a framework to study the general principles of regulatory relationships among enhancers in gene regulation.

## Results

### eNet builds enhancer networks based on single cell multi-omics data

Many enhancers exist as clusters in the genome; however, the underlying mechanism through which the clustered enhancers work together to regulate the same target gene remains largely unknown. To this end, we developed an algorithm eNet to build an enhancer network for each gene to quantitatively assess how multiple enhancers work together to regulate gene expression based on scATAC-seq and scRNA-seq data (**Methods**). The enhancer network we proposed is a gene regulation model that not only delineates the mapping between enhancers and target genes, but also quantifies the underlying regulatory relationships among enhancers, which differs from previous studies (Blobel et al., 2021; Hnisz et al., 2013; Ma et al., 2020; Osterwalder et al., 2018). First, given the scATAC-seq and scRNA-seq profiles, the enhancer accessibility and gene expression matrix of single cells were prepared as the input of eNet (**Figure 1A**). Second, a set of enhancers were identified, termed a putative enhancer cluster hereafter, which putatively regulate a specific target gene within a ±100 kb window based on the correlation between gene expression and enhancer accessibility in single-cell data (**Figure 1B**). Third, we evaluated the enhancer interaction potential based on their chromatin co-accessibility calculated by Cicero (Pliner et al., 2018), and determined the enhancer pairs with significantly high co-accessibility as the predicted enhancer interactions (PEIs) (**Figure 1C**). Fourth, an enhancer network was built to delineate how multiple enhancers interact with each other to regulate gene expression, where nodes represent enhancers and edges represent the PEIs between enhancers in a putative enhancer cluster (**Figure 1D**). Fifth, the complexity of the enhancer network was evaluated by two metrics: 1) the number of enhancers, termed the network size (*x*-axis), and 2) the frequency of PEIs, termed the network connectivity (*y*-axis), quantified by the average degree of network (Barabasi, 2016) (**Figure 1E**). Lastly, based on the network size and network connectivity, we classified the enhancer networks into several modes: Simple, Multiple, Complex and others (but will not be discussed due to limited cases) (**Figure 1F**). Intuitively, the complexity of the enhancer network increased from Simple mode to Multiple mode by involving more enhancers and further to Complex mode by increasing the interactions between enhancers. Altogether, eNet builds enhancer networks to clarify how a putative enhancer cluster regulates gene expression based on scATAC-seq and scRNA-seq data.

**Figure 1.**
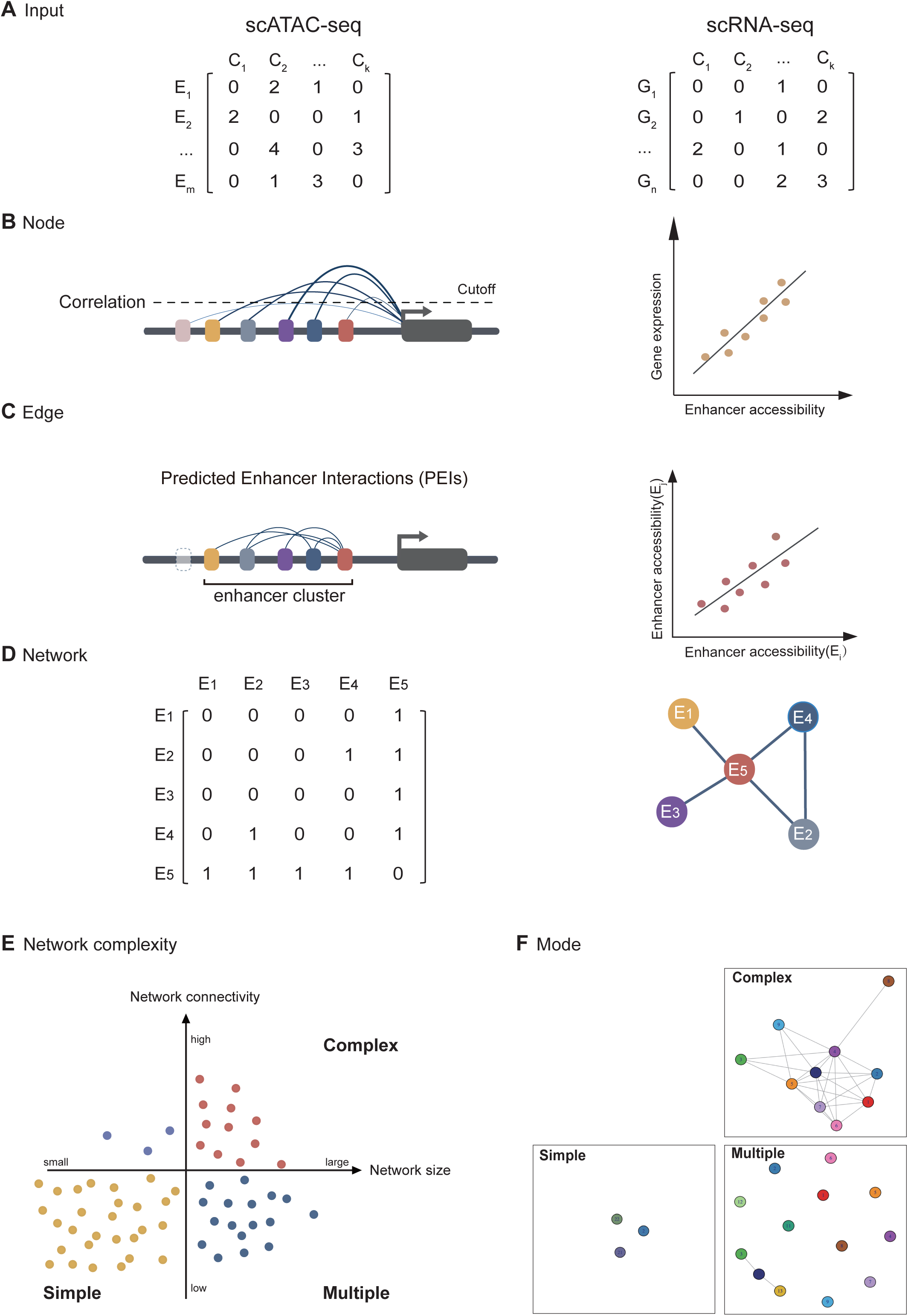
eNet, an algorithm to build enhancer networks based on scATAC-seq and scRNA-seq data. **(A)** Input: Preparation of the enhancer accessibility and gene expression matrix from scATAC-seq and scRNA-seq data. Each row represents an enhancer or a gene, while each column represents a cell. **(B)** Node: Identification of putative enhancer clusters regulating a specific target gene based on the correlation between gene expression and enhancer accessibility. **(C)** Edge: Determination of the predicted enhancer interactions (PEIs), the enhancer pairs with significantly high co-accessibility calculated using Cicero. **(D)** Network: Construct enhancer network to represent the PEIs among enhancers in a putative enhancer cluster, where nodes represent enhancers and edges represent PEIs. **(E)** Network complexity: Calculation of the network complexity by 1) network size, the number of enhancers (x-axis); and 2) network connectivity, the PEIs frequency, quantified by the average degree of network (y-axis). **(F)** Mode: Classification of the enhancer networks into three modes based on network complexity: Complex, Multiple and Simple, with representative examples shown in the cartoon.

### Cell identity and disease genes tend to be regulated by complex enhancer networks during human hematopoiesis

We first applied eNet to build enhancer networks during human hematopoiesis using a human blood dataset (Granja et al., 2019), including the single cell chromatin accessibility and transcriptional landscapes in human bone marrow and peripheral blood mononuclear cells (**Figure 2A**). In total, we built 11,438 enhancer networks during human hematopoiesis (**Figure 2B**). The number of enhancers in enhancer networks ranged from 1 to 50, with a median of 4 (**Figure S1A**). We noticed several blood-related cell identity or disease genes, such as *BCL11B*, *ETS1*, *CCR7* and *IL7R* displayed obviously large network size and high network connectivity (**Figure 2B**). This inspired the question that whether cell identity genes tend to be regulated by complex enhancer networks. To test this hypothesis, we classified these enhancer networks into three modes: Simple (controlled by one or few enhancers), Multiple (multiple enhancers but limited PEIs), and Complex (multiple enhancers and frequent PEIs). It resulted in 6,894 Simple, 2,992 Multiple and 1,552 Complex enhancer networks (**Figure 2B, Table S2 and Methods**). For example, the *CD3E* gene, encoding a subunit of the T-cell receptor-CD3 complex, was controlled by an enhancer network consisting of 14 PEIs among 9 enhancers (**Figure 2C**). In contrast, the *SERPINE2* gene, encoding a member of the serpin family of proteins that inhibit serine proteases, was controlled by an enhancer network containing the same number of enhancers but only 2 PEIs. Interestingly, the *CD3E* enhancer network showed significant higher chromatin co-accessibility than *SERPINE2*, irrespective of their indistinguishable chromatin accessibility and similar enhancer number (**Figure S1B** and **S1C**).

**Figure 2.**
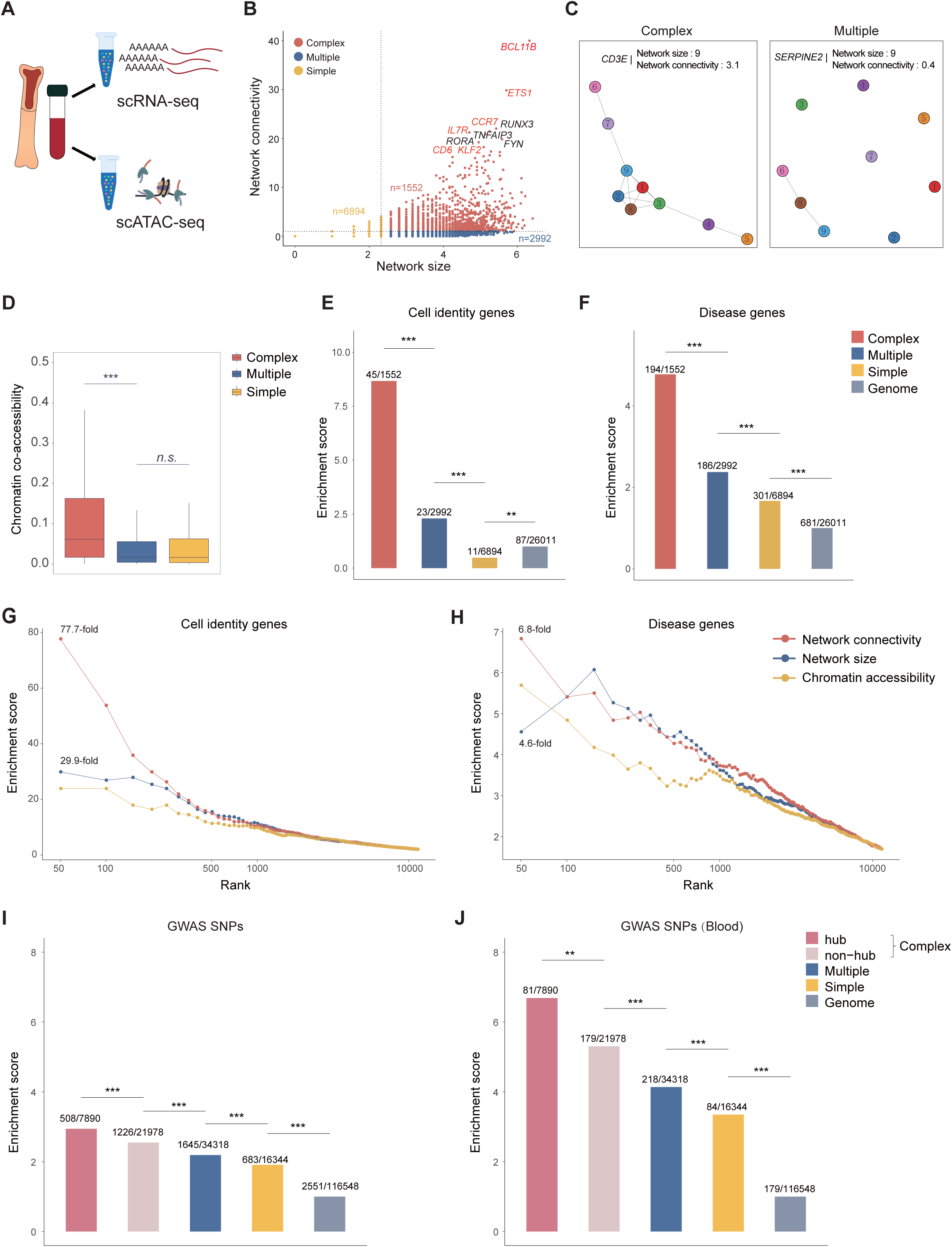
Enhancer networks during human hematopoiesis. **(A)** The human blood dataset. **(B)** Scatter plot of the enhancer networks during hematopoiesis, where the x-axis represents the network size (log_2_-scaled) and the y-axis represents network connectivity. Top 10 genes ranked by network connectivity are labelled, where known blood-related cell identity or disease genes are red-highlighted. **(C)** Representative enhancer networks in Complex or Multiple mode. **(D)** Chromatin co-accessibility of predicted enhancer interactions (PEIs) calculated using Cicero in Complex, Multiple and Simple modes. *p-*values were calculated using the Student’s *t*-test. *\*p* < 0.05*; **p* < 0.01*; ***p* < 0.001; *n.s.,* not significant. **(E** and **F)** Enrichment of cell identity (**E**) and disease genes (**F**) in genes in Complex, Multiple and Simple modes, using the whole genome as the background. The number of cell identity or disease genes and total genes in each group are labelled on each bar. *p*-values were calculated using the binomial test. *\*p* < 0.05*; **p* < 0.01*; ***p* < 0.001; *n.s.,* not significant. **(G** and **H)** Enrichment of cell identity (**G**) and disease genes (**H**) (y-axis) is plotted for top genes (x-axis) ranked by different properties of enhancer networks, including network connectivity (the frequency of PEIs in this study), network size (equivalent to the enhancer number in multiple enhancers (Osterwalder et al., 2018), DORCs (Ma et al., 2020)), or overall chromatin accessibility of enhancers (similar to the sum of the individual constituent enhancers in super-enhancers (Hnisz et al., 2013)). **(I)** Enrichment of the diseases/traits-related SNPs curated in the GWAS catalog for enhancers in Complex (hub and non-hub), Multiple, and Simple modes, using randomly selected genomic regions as the control. *p*-values were calculated using the binomial test. *\*p* < 0.05*; **p* < 0.01*; ***p* < 0.001; *n.s.,* not significant. **(J)** Enrichment of blood related GWAS SNPs.

Next, we curated a list of known cell identity genes in the blood system (**Methods, Table S3**) and calculated their enrichment in the genes regulated by three enhancer network modes (**Figure 2E**). We observed that genes regulated by Multiple mode showed higher enrichment in cell identity genes than those by Simple mode, which is consistent with previous reports that developmentally expressed genes are commonly associated with multiple enhancers (Ma et al., 2020; Osterwalder et al., 2018; Tsai et al., 2019). In addition, we found that genes regulated by Complex mode exhibited the highest enrichment in cell identity genes, 8.7-fold using the whole genome as the background (**Figure 2E**). Similarly, genes regulated by Complex mode displayed a higher enrichment in blood-related disease genes curated from DisGeNET (Pinero et al., 2017) than those by Multiple mode (4.8-fold vs. 2.4-fold, *p* = 3.4E-20, binomial test, **Figure 2F**). Notably, these observations were robust to various threshold values of network size and network connectivity (**Figure S2**). We also clarified that Complex mode enhancer network not mainly represents the enhancer networks with a stronger chromatin accessibility or larger enhancer number (network size) (**Figure S3 andS4**). These results suggested that cell identity and disease genes tend to be regulated by complex enhancer networks.

### Complexity of enhancer networks predicts cell identity and disease genes

To systematically evaluate the performance of the complexity of enhancer networks in predicting cell identity and disease genes, we ranked enhancer networks by the properties of enhancer networks, including network size, network connectivity, and overall chromatin accessibility. We then calculated the enrichment of cell identity and disease genes in the list of top ranked enhancer networks related genes, using the whole genome as the background (**Figure 2G** and **2H**). We found that the genes controlled by enhancer networks with more enhancers were overall preferentially more enriched for cell identity genes (**Figure 2G**), which concurs with previous studies (Hnisz et al., 2013; Ma et al., 2020; Osterwalder et al., 2018). Meanwhile, we observed an obvious correlation between network connectivity and the enrichment of cell identity genes. Importantly, network connectivity displayed better performance in predicting cell identity genes than the network size. For example, the top 50 genes ranked by network connectivity were 77.7-fold enriched of cell identity genes, compared with those by network size (29.9-fold), using the whole genome as the background. Both the network connectivity and network size showed remarkably better performance in predicting cell identity genes than the chromatin accessibility of enhancers in the network. Similarly, network connectivity displayed the best performance in predicting blood-related disease genes (6.8-fold in the top 50 genes, **Figure 2H**). Therefore, these analyses suggest complexity of enhancer networks can predict cell identity and disease genes.

### Network hub enhancers are functionally important

Enhancer networks provide an opportunity to study how individual elements work and then how they interact with each other to control gene expression. Toward this end, we focused on the enhancers with frequent PEIs in enhancer networks in Complex mode, termed network hub enhancers (**Methods**). To gain insight into the function of network hub enhancers in enhancer networks, we first compared the phastCons conservation scores (Siepel et al., 2005) of enhancers in Complex (hub and non-hub) and found that network hub enhancers displayed significantly higher level of sequence conservation than non-hub enhancers (*p* = 3.8E-8, Student’s *t*-test, **Figure S1D**), suggesting network hub enhancers might be more functionally important. Next, we assessed the enrichment of single-nucleotide polymorphisms (SNPs) linked to diverse phenotypic traits and diseases in the genome-wide association study (GWAS) catalog (Welter et al., 2014), in enhancers in Complex (hub and non-hub), Multiple, and Simple modes. We observed significantly higher enrichment of blood-associated GWAS SNPs in enhancers in Multiple mode than those in Simple mode (*p* = 2.8E-4, binomial test, **Figure 2I** and **2J**), which is consistent with previous studies (Hnisz et al., 2013; Osterwalder et al., 2018). Additionally, the enhancers in Complex mode (hub and non-hub) showed higher enrichment in GWAS SNPs associated with blood traits than those in Multiple mode. In particular, in Complex mode, hub enhancers displayed higher enrichment of GWAS SNPs associated with blood traits than non-hub enhancers (6.7-fold vs. 5.3-fold, *p* = 5.8E-3, binomial test, **Figure 2J**), suggesting hub enhancers might play important roles in enhancer networks. These results suggest that compared with Multiple and Simple modes, enhancers in Complex mode might be more important in diseases, where hub enhancers are major functional constituents.

### Enhancer network outperforms super-enhancer in predicting cell identity and disease genes

Super-enhancers (SEs) are clustered enhancers with a high density of transcriptional apparatus to drive robust expression of cell identity and disease genes (Hnisz et al., 2013). We next compared the performance of predicting cell identity and disease genes by enhancer networks and SEs. To this end, we downloaded a list of SEs associated with hematopoiesis-related cell types from the dbSUPER database (Khan and Zhang, 2016) and curated a catalog of hematopoiesis-related SEs containing 2,306 SEs (**Table S4**). We identified 2,159 potential target genes regulated by these SEs using ROSE algorithm (Hnisz et al., 2013). Comparing the genes regulated by SEs or by enhancer networks in Complex mode, we separated them into three groups: Complex-only (836), SE-only (1,443) and Complex SE (716) (**Figure 3A**). The constituent enhancers in these two groups (SE-only vs. Complex SE) showed significantly different chromatin co-accessibility, but indistinguishable chromatin accessibility (**Figure 3B** and **3C**). It might explain the diverse and heterogeneous mechanisms of SEs, such as cooperative, redundant and hierarchical revealed by CRISPR/Cas9 genome editing assays. Strikingly, genes in Complex-only group displayed significantly higher enrichment in cell identity and disease genes than those in SE-only group, while genes in Complex SE group showed the highest enrichment (**Figure 3D** and **3E)**. Moreover, we observed similar patterns in GM12878 cell line that enhancer networks precede SEs in predicting cell identity and disease genes (**Figure S5A**-**S5C)**. We further ranked genes by network connectivity, network size, chromatin accessibility or SE ranks based on H3K27ac signal, and found that network connectivity showed the best performance in predicting both cell identity and disease genes, comparing with network size, chromatin accessibility and SE ranks (**Figure 3F** and **3G**). These results suggested that the enhancer networks precede SEs in predicting cell identity and disease genes.

**Figure 3.**
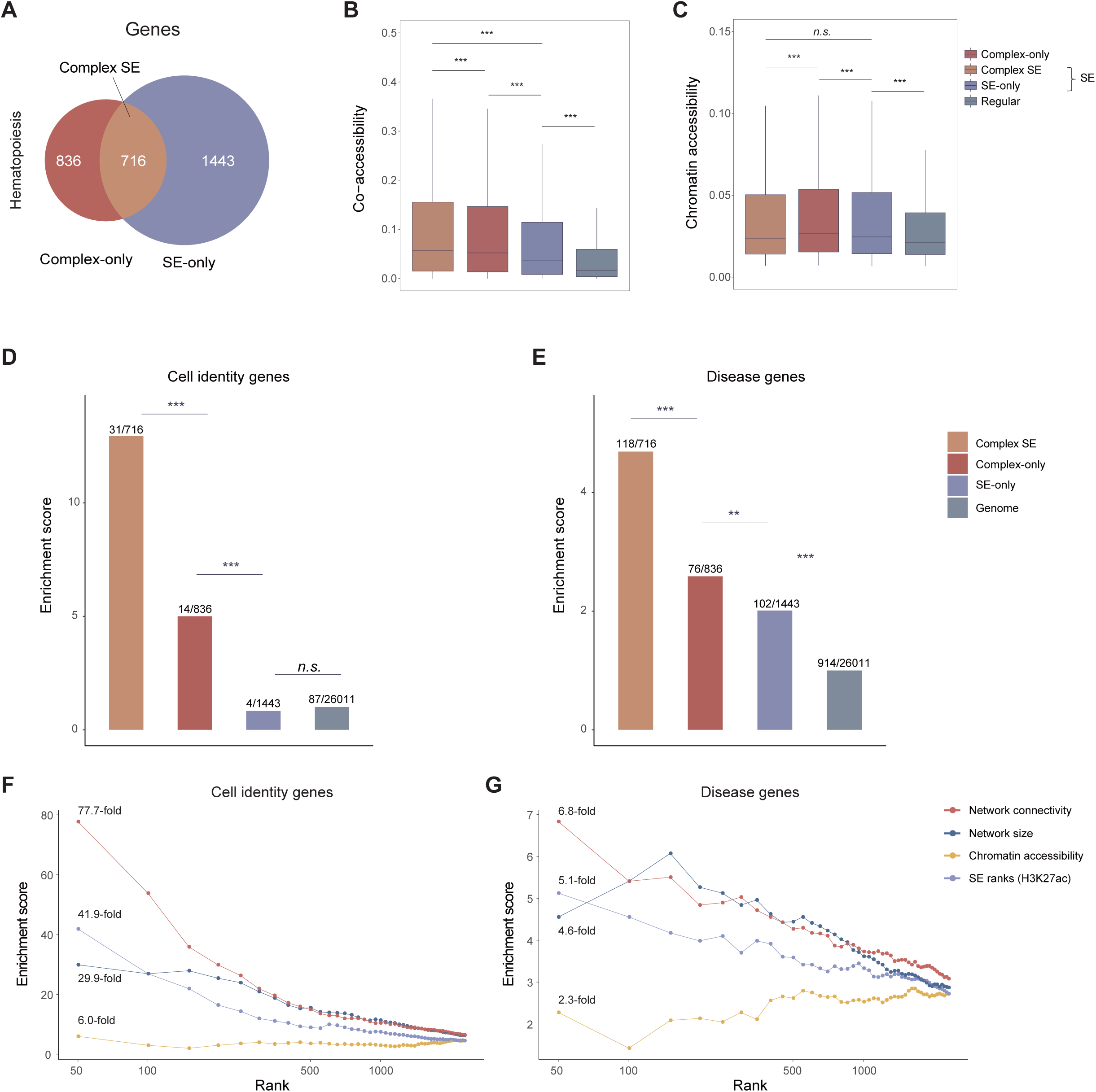
Enhancer network outperforms super-enhancer in predicting cell identity and disease genes. **(A)** Venn diagram showing the overlap between genes in Complex mode in Figure 2 in blood dataset and hematopoiesis-related SEs, resulting in three groups, Complex-only, Complex SE (SEs with network structure) and SE-only (SEs without network structure) **(B, C)** Co-accessibility (**B**) and chromatin accessibility (**C**) of the constituent enhancers in three groups, using regular enhancers as control. p-values were calculated using Student’s t-test. \**p* < 0.05; \*\**p* < 0.01; \*\*\**p* < 0.001; *n.s.*, not significant. **(D** and **E)** Enrichment of cell identity (**D**) and disease genes (**E**) in genes in three groups, using the whole genome as the background. *p-*values were calculated using the binomial test. *\*p* < 0.05*; **p* < 0.01*; ***p* < 0.001; *n.s.,* not significant. **(F** and **G)** Enrichment of cell identity (**F**) and disease genes (**G**) (y-axis) for the genes in (**A**), ranked by the network complexity (x-axis), measured by 1) network connectivity, as well as the overall enhancer activity, 2) network size as well as enhancer number, 3) chromatin accessibility and 4) SE ranks based on H3K27ac signals calculated by ROSE (Hnisz et al., 2013).

### Enhancer networks based on PEIs remedy the resolution limitations in Hi-C chromatin interactions

The proximity ligation-based methods to capture genome-wide chromatin interactions at high-resolution for the analysis of enhancer interactions remains difficult and costly (Lieberman-Aiden et al., 2009; Mumbach et al., 2017; Tang et al., 2015). We wonder to what extent the PEIs in eNet analysis resolve the resolution limitations in Hi-C data. To this end, we compared enhancer networks based on PEIs and Hi-C data in GM12878 cell line (human B-lymphoblastoid cells), where scATAC-seq (Ma et al., 2020), H3K27ac ChIP-seq (Consortium, 2012) and high-resolution Hi-C data (Rao et al., 2014) are available. We observed the high co-accessible enhancer pairs (PEIs) showed significant enrichment of Hi-C chromatin interactions (**Figure 4A**), indicating the overall concordance between co-accessible pairs and proximity ligation-based chromatin interactions (Pliner et al., 2018). For example, at the locus controlling *CCR7*, a gene expressed in various lymphoid tissues and activates B and T lymphocytes, we predicted 20 PEIs based on scATAC-seq data, while only 10 chromatin interactions were detected via Hi-C probably due to the limited resolution at 5kb level (**Figure 4B-C**). We systematically compared the enhancer networks based on scATAC-seq and Hi-C data by replacing PEIs with Hi-C interactions and re-built enhancer networks. We observed a significant overlap between the genes controlled by the complex enhancer networks based on PEIs and Hi-C data (**Figure 4D,** *p* < 2.2E-16, Fisher’s exact test). Interestingly, PEI-only group showed significant higher enrichment of cell identity and disease genes than HiC-only group, where PEI-with-HiC showed the highest enrichment (**Figure 4E** and **4F**). Moreover, we found the network hub enhancers derived from PEIs showed significant higher enrichment of GWAS SNPs than those from Hi-C data (**Figure S5D**-**S5F**). Taken together, these results suggested enhancer networks based on PEIs remedy the resolution limitations of chromatin interactions in Hi-C data.

**Figure 4.**
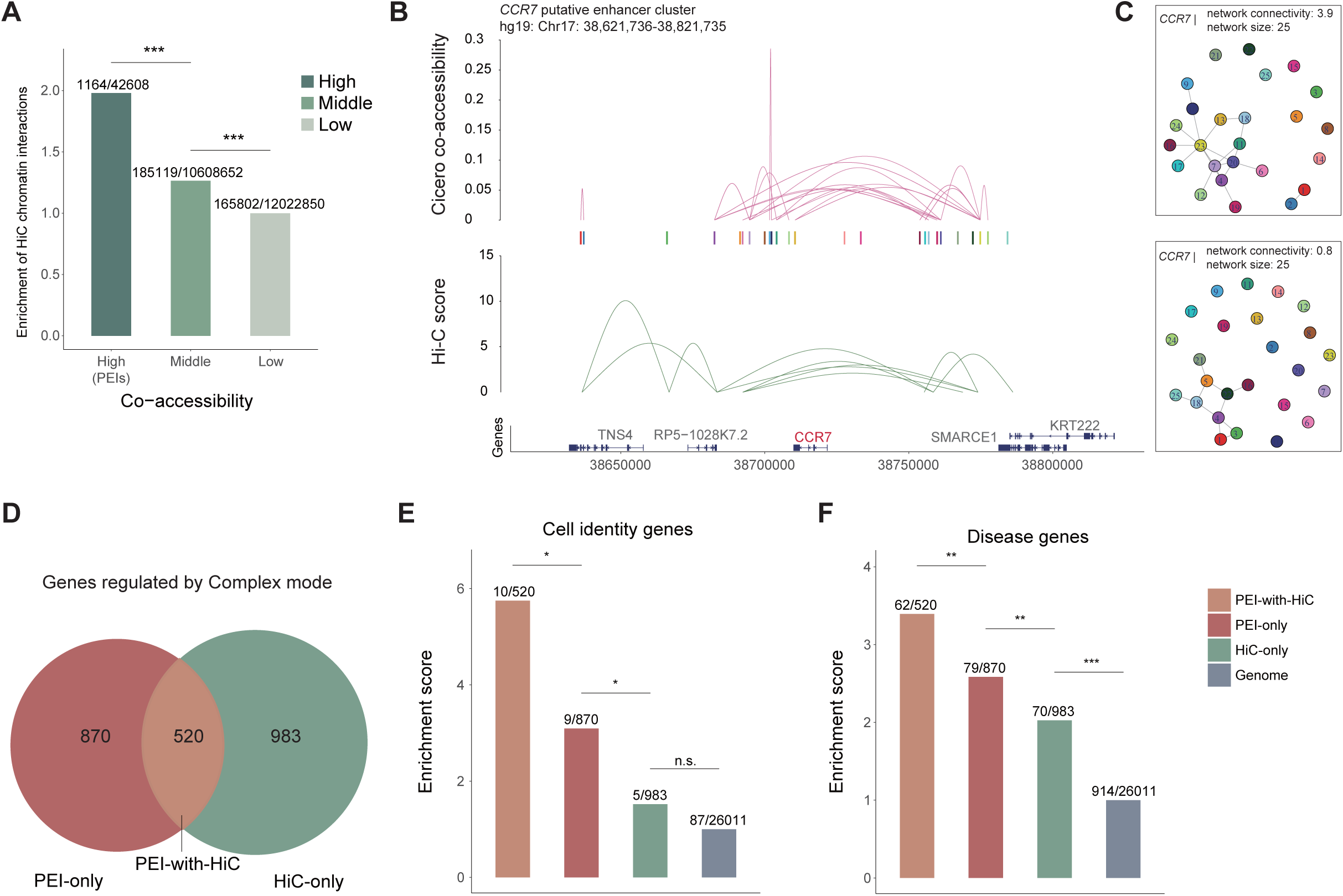
Comparison of enhancer networks based on PEIs and Hi-C chromatin interactions in GM12878 cell line. (**A**) Enrichment of chromatin interactions detected by Hi-C in three groups of enhancer pairs ranked by chromatin co-accessibility: High (PEIs), Middle and Low, using the group Low as the background. p-values were calculated using binomial test. \**p* < 0.05; \*\**p* < 0.01; \*\*\**p* < 0.001; *n.s.*, not significant. (**B**) Cicero connections for the *CCR7* locus compared to Hi-C (Rao et al., 2014). Link heights for Hi-C are the interaction frequency of each chromatin interaction. (**C**) *CCR7* enhancer networks built based on PEIs (above) or Hi-C chromatin interactions (bottom). (**D**) Venn diagram showing the overlap of genes regulated by the Complex enhancer networks defined based on PEIs and Hi-C data, resulting in three groups: PEIs-with-HiC, PEIs-only, and HiC-only. (**E** and **F**) Enrichment of cell identity (**E**) and disease genes (**F**) in three groups, using the whole genome as the background. p-values were calculated using the binomial test. \**p* < 0.05; \*\**p* < 0.01; \*\*\**p* < 0.001; *n.s.*, not significant.

### Dynamics of PAX5 enhancer network drives gene expression during B cell lineage commitment

Enhancer networks were built based on single cell multi-omics data, providing an opportunity to investigate the dynamic role of enhancer networks in determining gene expression during cell differentiation. To this end, we focused on B cell differentiation, from hematopoietic stem cell (HSC), lymphoid-primed multipotent progenitor (LMPP), common lymphoid progenitor (CLP), pre-B, to B cells (**Figure 5A** and **Methods**). The *PAX5* gene, a known key regulator for B cell differentiation, specifically expressed in pre-B and B cells, was controlled by a putative enhancer cluster consisting of 24 enhancers (**Figure 5B** and **5C**). To understand the relationship between these constituent enhancers and their roles in regulating gene expression during cell differentiation, we built cell-type-specific enhancer networks by constructing the enhancer networks for each cell type independently (**Methods**). Comparing the enhancer networks specific for HSC, LMPP, CLP, pre-B, and B cells, we observed the sequential changes in the constituent enhancers during B cell differentiation, in terms of both chromatin accessibility and network interactions (**Figure 5D**). Within the *PAX5* enhancer network, we noticed that enhancer E14, constitutively accessible from HSC to B cells, functions as a network hub enhancer to coordinate enhancer network interactions to establish the enhancer network gradually during B cell differentiation (**Figure 5D**). Interestingly, we found that the *PAX5* enhancer network was almost fully established in the CLP and pre-B stages, which preceded the gene expression of *PAX5* in pre-B and B cells (**Figure 5C** and **5D**). It suggests the establishment of an enhancer network may drive gene expression during lineage commitment.

**Figure 5.**
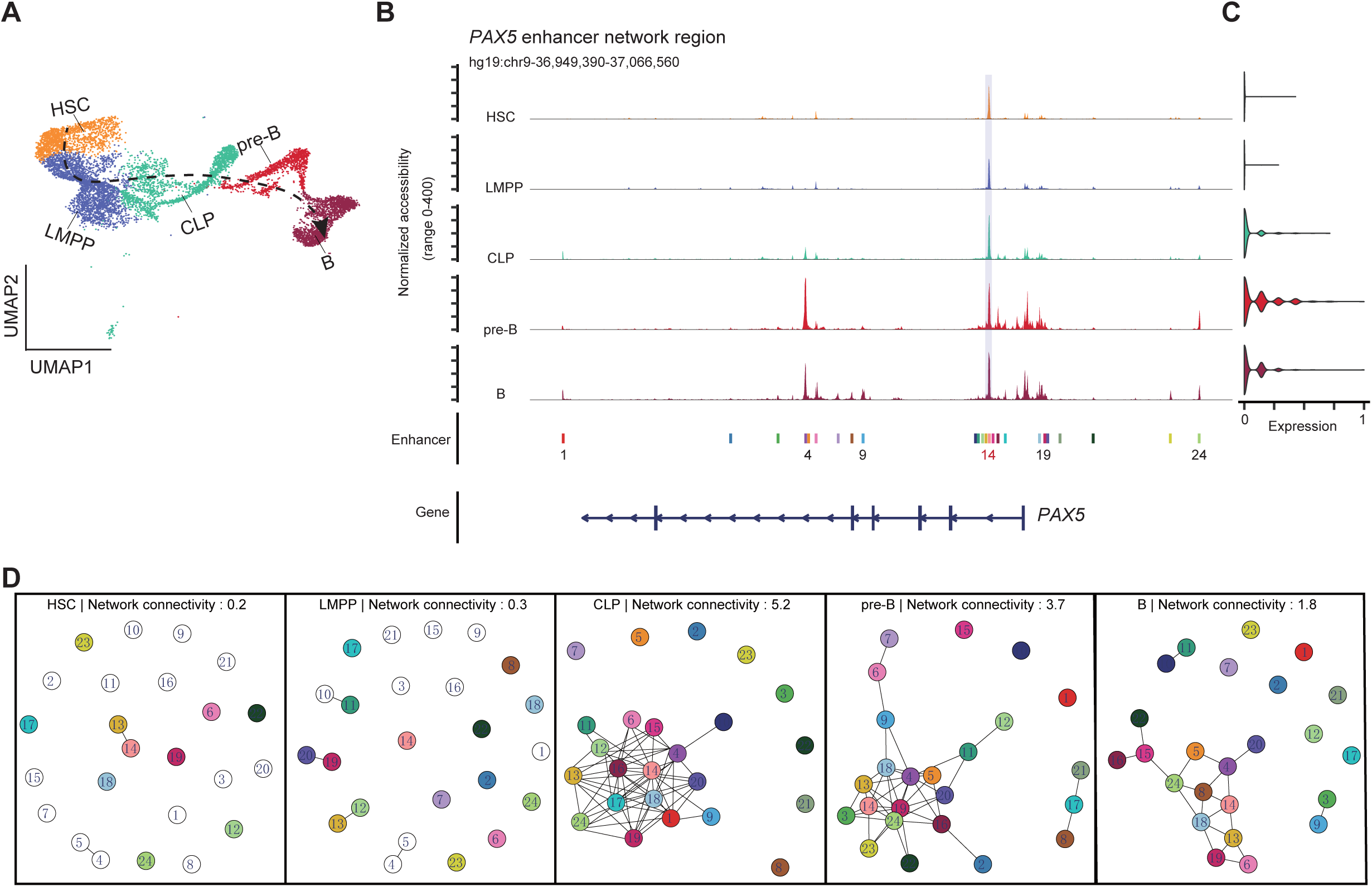
Dynamics of enhancer networks during B cell differentiation. **(A)** UMAP of B cell differentiation colored by cell type annotation, the dash-line indicates the pseudotime during B cell differentiation inferred based on scATAC-seq data. **(B)** Genome browser track of *PAX5* putative enhancer cluster (*n*=24) that are accessible at any one of the developmental stages of HSC, LMPP, CLP, pre-B, and B cell types. **(C)** Violin plot showing *PAX5* expression. **(D)** The *PAX5* enhancer networks in HSC, LMPP, CLP, pre-B, and B cells, where the colored nodes represent accessible enhancers while the empty nodes represent closed enhancers. The edges represent PEIs.

To test this hypothesis, we performed trajectory analysis for B cell differentiation using the method described before (Satpathy et al., 2019) to order cells in pseudotime based on scATAC-seq data in HSC, LMPP, CLP, pre-B, and B cells (**Figure S5G**). We then systematically compared gene expression, chromatin accessibility, and enhancer network connectivity along the B cell differentiation pseudotime (**Figure S5H**). We quantified the differences in pseudotime of B cell differentiation between the onset of gene expression and establishment of the enhancer network (**Methods**). Notably, there was a lag of pseudotime between the onset of gene expression and chromatin accessibility (*p* = 2.8E-2, Student’s *t*-test, **Figure S5H** and **S5I**), which supports the hypothesis that chromatin accessibility is a marker for lineage-priming (Lara-Astiaso et al., 2014; Ma et al., 2020; Rada-Iglesias et al., 2011). More importantly, we found enhancer networks were established earlier than gene expression occurred (*p* = 2.9E-6, Student’s *t*-test, **Figure S5H** and **S5I**), even prior to the change in chromatin accessibility, suggesting the dynamics of enhancer networks drive gene expression during cell differentiation. Taken together, we demonstrate that enhancer networks are established gradually during lineage commitment, which drives the expression of cell identity genes.

### eNet is robust and broadly applicable

To investigate the broad applicability of eNet, we applied it to various datasets in human or mouse tissues across different single-cell platforms, including SHARE-seq mouse skin dataset (Ma et al., 2020), SNARE-seq mouse cerebral cortex dataset (Chen et al., 2019) and sci-ATAC-seq3 human fetal kidney and heart datasets (Domcke et al., 2020). Similar to the above findings, we found cell identity and disease genes tended to be regulated by complex enhancer networks (**Figure S6A**, **S6C**, **S6E**, and **S6G**). The network connectivity showed the best performance in predicting cell identity genes and disease genes (**Figure 6A, 6C, 6E, 6G, S6B, S6D, S6F** and **S6H**). Hub enhancers in Complex mode displayed the highest enrichment of tissue-related GWAS SNPs (**Figure 6B**, **6D**, **6F** and **6H**). These analyses in various human or mouse tissues datasets (**Figure 2**, **6** and **S6**) support the conclusion that eNet is robust and broadly applicable in various biological systems and different single-cell platforms.

**Figure 6.**
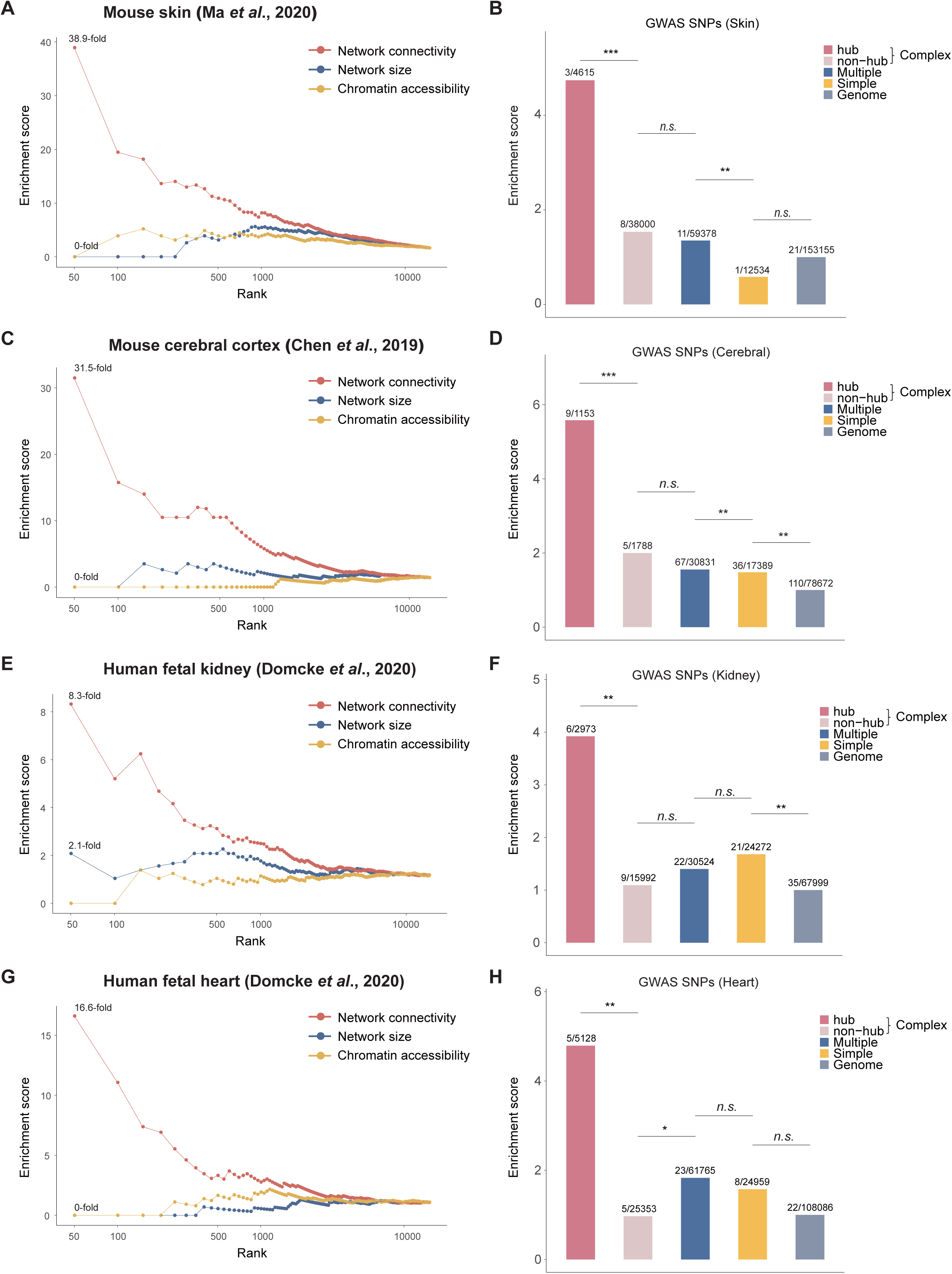
Enhancer networks in various human or mouse tissues across different single-cell platforms. **(A, C, E, G)** Enrichment of cell identity genes (y-axis) is plotted for top genes ranked by various scoring methods (x-axis) in different tissues and approaches. (**A**) mouse skin dataset (SHARE-seq) (Ma et al., 2020), (**C**) mouse cerebral cortex dataset (SNARE-seq) (Chen et al., 2019), (**E**) human fetal kidney dataset (sci-ATAC-seq3) (Domcke et al., 2020), and (**G**) human fetal heart dataset (sci-ATAC-seq3) (Domcke et al., 2020). **(B, D, F, H)** Enrichment of tissue-related diseases/traits SNPs curated in GWAS catalog in enhancers in Complex (hub and non-hub), Multiple, and Simple modes, using randomly selected genomic regions as the control. (**B**) mouse skin dataset, (**D**) mouse cerebral cortex dataset, (**F**) human fetal kidney dataset, and (**H**) human fetal heart dataset. *p-*values were calculated using the binomial test. *\*p* < 0.05*; **p* < 0.01*; ***p* < 0.001; *n.s.,* not significant.

### Model of enhancer networks in gene regulation

Our analysis revealed three modes of enhancer networks in regulating gene expression according to their network complexity: Complex, Multiple, and Simple. To further understand the underlying biological functions and mechanisms, we evaluated the functional enrichment of genes regulated by these three modes (**Figure 7A**). We found genes regulated by the Simple mode were primarily enriched in housekeeping functions, such as RNA modification and DNA repair (**Figure 7A**). In contrast, genes regulated by the Complex mode were enriched in key genes related to cell fate commitment, such as the regulation of leukocyte differentiation in human blood, skin development in mouse skin and cerebellar cortex formation in mouse cerebral datasets. Meanwhile, genes in Multiple mode were enriched in a mixture of both housekeeping and cell identity functions. In addition, Complex mode preferentially regulated upstream regulators, such as transcription factors (Lambert et al., 2018), which was observed in all three datasets (**Figure S7A**).

**Figure 7:**
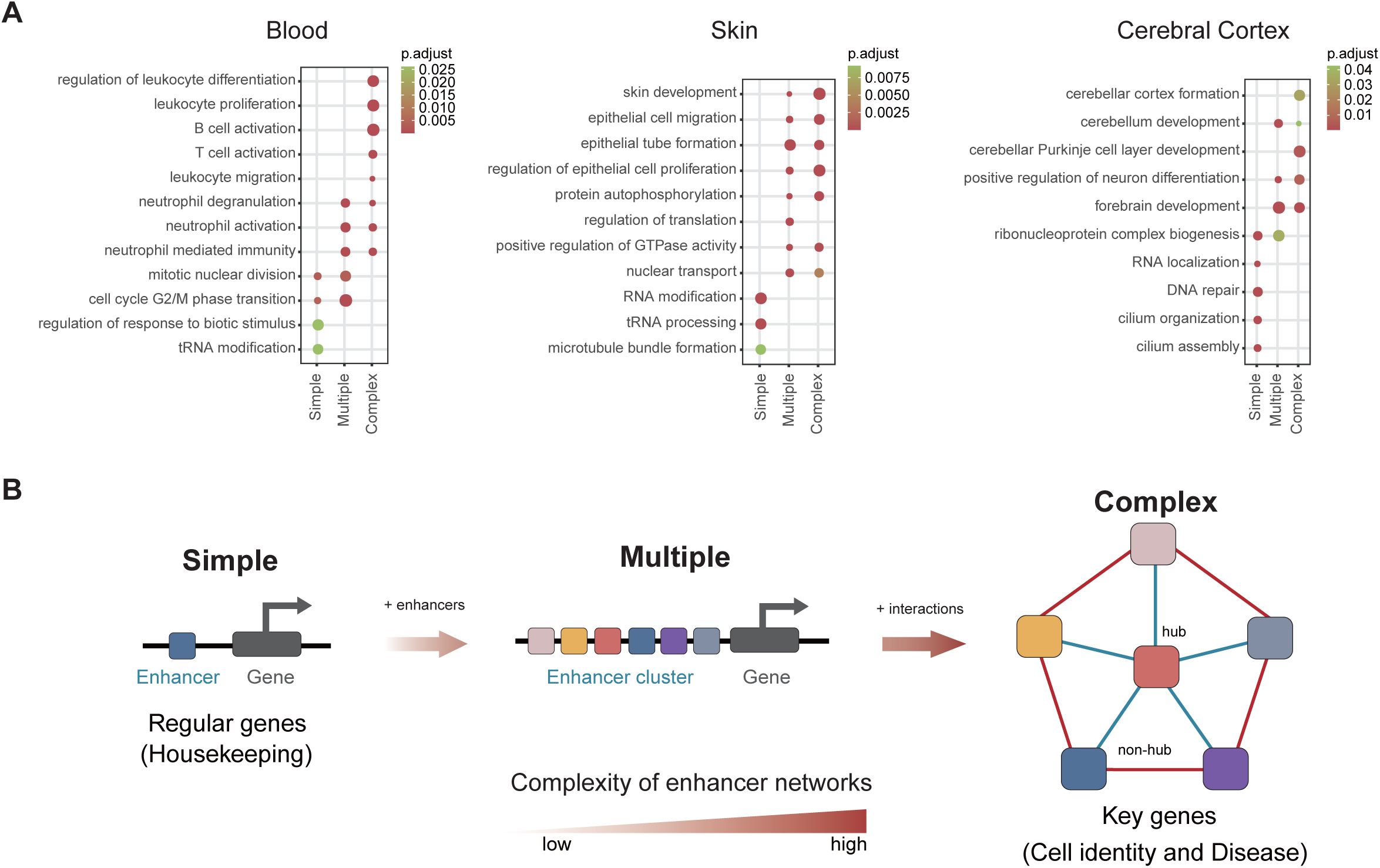
Model of enhancer networks in gene regulation. **(A)** Functional enrichment of genes regulated by enhancer networks in Simple, Multiple, and Complex modes in human blood, mouse skin, and mouse cerebral cortex datasets. **(B)** Three modes of enhancer networks. Simple mode, involving one or very few enhancers, provides quick response to control a large amount of regular genes, such as housekeeping genes, at low cost; Multiple mode, involving multiple enhancers but limited PEIs, increases regulation strength as well as redundancy at the cost of the number of enhancers (nodes); Complex mode, involving multiple enhancers and frequent PEIs, provides robustness of gene regulation for key genes, such as cell identity and disease genes, at the cost of edges, where hub enhancers are functionally important.

Therefore, we proposed a model of enhancer networks containing three modes according to their network complexity: Simple, Multiple, and Complex (**Figure 7B**). By definition, in Simple mode, gene regulation was controlled simply by one or a limited number of enhancers; we speculated it provided a quick response to control a large amount of regular genes, such as housekeeping genes, at a low cost. Meanwhile, in Multiple mode, gene regulation was controlled by multiple enhancers but limited PEIs; this might increase the strength of regulation and redundancy of gene expression at the cost of involving more enhancers. Lastly, gene regulation was controlled by multiple enhancers and frequent PEIs in Complex mode, perhaps the most robust to random failures of individual enhancers (transcriptional noise or genetic mutation), at the cost of connecting enhancers and primarily controls key cell identity genes. Enhancer networks are established gradually during lineage commitment and drive the expression of cell identity genes, where network hub enhancers play central roles to coordinate the network system.

## Discussion

### The concept of complexity of enhancer networks

The term ‘enhancer’ first appeared to describe the effects of SV40 DNA sequences on the expression of a β-globin gene (Banerji et al., 1981). Since then, hundreds of thousands of enhancers have been nominated via genome-wide biochemical annotations (Gasperini et al., 2020; Neph et al., 2012). However, only a small number of enhancers have been functionally tested. While multiple enhancers existing as clusters in the genome to regulate the same target gene is prevalent, which are known to provide phenotypic robustness in development (Osterwalder et al., 2018; Perry et al., 2011), the underlying mechanisms remain largely unknown.

Enhancer networks have been reported in previous studies. For example, Malin et al. constructed enhancer network based on the correlated DNase hypersensitivity of enhancers across 72 cell types (Malin et al., 2013). Chen et al. built tissue-specific enhancer functional networks for associating distal regulatory regions to disease by integrating thousands of epigenetics and functional genomics data sets (Chen et al., 2021). Carleton et al. discovered the enhancer combinations by targeting a set of 10 enhancers by simultaneous deactivation of multiple enhancers using CRISPR-based technique (Carleton et al., 2017). However, it is infeasible to scale up this approach to rigorously test a wide range of enhancers due to technical difficulties. Proximity ligation-based methods, including ChIA-PET and Hi-C (Lieberman-Aiden et al., 2009; Mumbach et al., 2017; Tang et al., 2015), capture chromatin interactions, but limited to the resolution. Most of ChIA-PET or Hi-C data can only achieve a resolution at 5-20kb, which is not sophisticate enough for enhancer studies at ∼500bp (the median of enhancer length). In this study, we reported an algorithm eNet to build enhancer network per gene based on the rich source of single-cell multi-omics data and greatly extended these previous findings in understanding the biological relevance and implications of enhancer network. Most importantly, to our knowledge, we for the first time propose the concept of “complexity of enhancer network” and establish its functional links with cell identity or disease. Furthermore, our study overcomes the above limitations on resolution and scalability by integrating single-cell multiple omics data to quantify enhancer interactions.

Chromatin co-accessibility based on scATAC-seq data has been used before, but mainly for connecting enhancers to their putative target genes (Pliner et al., 2018). We quantified the complexity of the enhancer network by two metrics: network size and network connectivity. The first metric, network size (the number of enhancers) is equivalent or similar to the sum of the individual constituent enhancers in an enhancer cluster reported in previous studies, such as multiple enhancers (Osterwalder et al., 2018), domains of regulatory chromatin (DORCs) (Ma et al., 2020), regulatory locus complexity (Gonzalez et al., 2015) or super-enhancers (Hnisz et al., 2013). However, the second metric, network connectivity (the frequency of PEIs), measuring the potential enhancer interactions based on their chromatin co-accessibility, differs from these existing studies. Thus, the enhancer network not only delineates the mapping between enhancers and the target gene, but also clarifies the underlying regulatory relationship between enhancers. We applied eNet in various biological systems and found the number of enhancers in the network was correlated with the importance of target genes, which was expected and consistent with previous studies (Gonzalez et al., 2015; Hnisz et al., 2013; Ma et al., 2020; Osterwalder et al., 2018). Strikingly, we further found that network connectivity had the best performance in predicting cell identity and disease genes, where the network hub enhancers are the most functionally important in the network. The enhancer networks concept might be also helpful to interpret the phase separation model for gene regulation (Sabari et al., 2018), *e.g.* whether the genes regulated by Complex mode are more likely to form the phase separation or whether network hub enhancers play a role in mediating the phase separation.

In network science, the hierarchical organization, or hub-and-spoke network, is robust to random failures, as only the failure of its central hub node can break the network into isolated components (Barabasi, 2016). However, it has a low tolerance to an attack that removes its central hub. Thus, we wondered whether connections between non-hub enhancers are dispensable in enhancer networks. To this end, we quantified the network connectivity in an alternative way by maximum degrees of nodes in the network, which represented the importance of the central hub node in a network, termed as network connectivity (maximum), instead of by average degrees of nodes in the network as described above, termed as network connectivity (average). Surprisingly, we observed that the performance of the network connectivity (maximum) markedly decreased in the prediction of both cell identity and disease genes, compared to network connectivity (average) (**Figure S7B** and **S7C**). It suggested that the connections between non-hub enhancers are also indispensable in the enhancer network, which further complements our previous model based on Hi-C chromatin interactions(Huang et al., 2018). We speculated that by connecting some of non-hub nodes, the reinforced network model has a higher tolerance to targeted attacks, where the removal of the hub does not fragment the network. In this way, enhancer networks provide the robustness in biological systems to random failures (e.g. transcriptional noise) as well as attack on the central network hub enhancers (e.g. genetic mutation). Taken together, the concept of complexity of enhancer networks allows us to identify key cell identity and disease genes and to explore the underlying mechanisms, such as how constituent enhancers interact with each other to regulate gene expression.

### Super-enhancers in gene regulation

Super-enhancers exhibit disproportionately higher signals for the enhancer marks (such as H3K27ac and binding of Mediator and TFs), which control cell identity and disease genes (Hnisz et al., 2013). While SEs have attracted enormous interest in further studying these interesting regulatory elements, it remains datable on how functionally and mechanistically distinct a super-enhancer is from a typical enhancer as they are defined operationally but not functionally (Blobel et al., 2021; Pott and Lieb, 2015). For example, the current dissection of individual enhancers suggests that the mechanistic relationships among constituent enhancers of SEs are highly diverse and heterogeneous, such as cooperative, redundant, hierarchical, or temporal (Bahr et al., 2018; Cai et al., 2020; Canver et al., 2015; Fulco et al., 2016; Hay et al., 2016; Huang et al., 2016; Kai et al., 2021; Shin et al., 2016). Here, we found SEs can be subdivided into two groups SE-only (SEs without network structure) and Complex SEs (SEs with network structure), which displayed different in chromatin co-accessibility between the constituent enhancers, irrespective of their indistinguishable chromatin accessibility. Thus, this distinct feature, with or without network structure, might explain their diverse and heterogeneous mechanisms. Furthermore, SEs are identified computationally by the linear clustering of individual components in the genome (Hnisz et al., 2013). This ignores the observation that a gene can be regulated by multiple enhancers which are not constrained by linear genome distances. In contrast, eNet assigns each individual enhancer to its potential target gene based on the correlation between gene expression and enhancer accessibility across various cells (**Figure 1B**). This approach identifies a more complete set of the enhancers that regulate a specific gene, irrespective of the distance between enhancers.

### Enhancer networks in single cells

Currently, most studies on enhancer clusters rely on chromatin marks and the binding of Mediator and TFs at bulk population (Hnisz et al., 2013). It remains largely unknown whether enhancer clusters control robust gene expression simply through population averaging. The development of scATAC-seq and scRNA-seq technologies generated a large amount of single cell multi-omics profiles in various biological systems (Argelaguet et al., 2019; Granja et al., 2019; Sarropoulos et al., 2021; Trevino et al., 2021) could be leveraged towards addressing this question. However, these studies have largely focused on connecting distal enhancers with their target genes, but rarely on exploring the underlying regulatory relationships among enhancers regulating the same gene, namely, the enhancer networks in this study. Here, as the primary feature distinguishing our work from these studies, eNet allows us to explore how individual elements interact with each other to control gene expression during lineage commitment at single-cell resolution, as illustrating by the example of *PAX5* enhancer network. As the second advantage, by building enhancer networks based on single cell multi-omics data, eNet is an unsupervised approach to simultaneously identify key cell identity and disease genes and the underlying enhancer regulatory relationships. Thus, it is not necessary to know the cell identity in advance from primary samples or conduct challenging experimental steps, such as cell subpopulation isolation and chromatin immunoprecipitation sequencing (ChIP-seq).

### Limitations

The primary limitation of our work is the lack of experimental validation on the regulatory role of enhancer networks during development and disease. In a parallel study, Shu et al. performed LacZ transgenic mouse assay and *in vivo* enhancer perturbation by CRISPR/Cas9-mediated genome editing and found that the network hub enhancers played a central role in orchestrating spatiotemporal gene expression programs of *Atoh1* during spinal cord development (also see “related manuscript file” for details). Fully determining the regulatory roles of enhancer networks requires more comprehensive investigations in future, such as combining epigenetic features, chromatin looping, reporter assays, and enhancer perturbations in relevant cell lines, and *in vivo* models. Moreover, one of the motivations of our study is that it currently remains difficult and costly to capture genome-wide chromatin interactions at high-resolution by proximity ligation-based methods for the analysis of enhancer interactions (Lieberman-Aiden et al., 2009; Mumbach et al., 2017; Tang et al., 2015). To address this question, eNet builds enhancer networks based on the assumption that the Cicero-detected significant co-accessible pairs (Pliner et al., 2018), the predicted enhancer interactions (PEIs) used in this study, are overall concordant with proximity ligation-based chromatin interactions. Analysis in GM12878 cell line revealed network hub enhancers overlapped with part of Hi-C hub enhancers. Meanwhile, it captured significant fraction of distinct enhancers, which were functionally important.

While it might be due to the limited resolution of current Hi-C data (Rao et al., 2014), it is also important to recognize that inconsistencies exist between these two measurements. Thus, it is important to systematically compare the coherence of the enhancer networks from scATAC-seq with those from proximity ligation-based chromatin interactions at higher resolution. In this sense, eNet can be easily applied to high-resolution chromatin interaction data, if available in the future.

## Materials and Methods

### Data Sources

The scATAC-seq and scRNA-seq datasets used in this study were obtained from the literature. The human blood dataset includes single cell profiling of gene expression and chromatin accessibility in human primary bone marrow and peripheral blood mononuclear cells measured by the Chromium platform (10x Genomics)(Granja et al., 2019). The mouse skin dataset contains the single cell profiling of gene expression and chromatin accessibility during mouse skin development measured by SHARE-seq(Ma et al., 2020). The mouse cerebral cortex dataset consists of the single cell profiling of gene expression and chromatin accessibility of developing mouse cerebral cortex measured by SNARE-seq(Chen et al., 2019). The human fetal kidney and heart datasets include single cell profiling of gene expression and chromatin accessibility of human fetal kidney and heart measured by sci-ATAC-seq3(Domcke et al., 2020) and sci-RNA-seq3(Cao et al., 2020).A list of all used datasets and accession numbers are summarized in **Table S1**.

### eNet

eNet is an algorithm to build enhancer networks for clustered enhancers controlling the same gene based on scATAC-seq and scRNA-seq datasets. Briefly, it contains the following six steps.

#### Step 1. Preparing input matrix (Input)

In this study, the processed single cell chromatin accessibility and gene expression matrix data were downloaded directly from public literatures and used as the input for eNet.

#### Step 2. Identifying the putative enhancer cluster (Node)

The chromatin accessible regions outside of ± 2 kb of transcriptional start sites (TSS) were considered enhancers. We identified a set of enhancers, as the nodes in the network, which putatively regulate a specific target gene based on the correlation between gene expression and enhancer accessibility across various cells by adapting the method previously described (Li et al., 2021; Ma et al., 2020), with some modifications. Briefly, given a gene, we first selected the enhancers located within a ± 100 kb window around each annotated TSS as enhancer candidates. For each gene-enhancer pair, we then calculated the Spearman correlation between enhancer chromatin accessibility and gene expression. The Spearman correlations were z-score normalized using genome-wide gene-enhancer pairs as the background. Lastly, by defining a cut-off at the z-score with an empirically defined significance threshold of *p*-value < 0.01 (one-sided Student’s *t*-test), we identified a putative enhancer cluster regulating the specific target gene.

#### Step 3. Identifying the predicted enhancer interactions (Edge)

We determined the potential chromatin interactions between enhancers within each putative enhancer cluster as the edges of the network. The chromatin co-accessibility of enhancer pairs across various cells was calculated using Cicero (Pliner et al., 2018), a method that predicts cis-regulatory DNA interactions from single-cell chromatin accessibility data. By applying a threshold value of the co-accessibility calculated, we determined the significant co-accessible enhancer pairs, termed as the predicted enhancer interactions (PEIs).

#### Step 4. Building enhancer networks (Network)

We built a binary adjacency matrix to represent the predicted enhancer interactions for each putative enhancer cluster, where 1 or 0 represent two enhancers with or without predicted enhancer interactions, respectively. Thus, the adjacency matrix can be visualized as an enhancer network, where nodes represent enhancers and the edges represent PEIs.

#### Step 5. Calculating network complexity (Network complexity)

We evaluated the complexity of the enhancer networks by the network size and connectivity. Network size was quantified by the quantity of nodes in the network. Network connectivity was quantified by the average degree (Barabasi, 2016), which were calculated as two-fold of the number of edges and divided by the number of nodes.

#### Step 6. Classification of enhancer networks (Mode)

We built the enhancer network for each gene genome-wide by repeating from steps 1-5. Then, by applying a threshold value of network size and connectivity, we can classify the enhancer networks into several groups: Complex (large size and high connectivity), Multiple (large size but low connectivity), Simple (small size and low connectivity) and others (small size but high connectivity, not discussed due to limited cases).

#### Defining network hub enhancers for enhancer networks in Complex mode

In Complex mode, we calculated the node degree for each enhancer and normalized them by the total number of edges in network, termed as normalized node degree. By applying a threshold value of the normalized node degree, we divided the enhancers into two groups, termed as network hub enhancers and non-hub enhancers, where network hub enhancers are those with high frequency of PEIs.

### Robustness analysis of eNet

#### Building weighted enhancer network in Step 4

In additional to the binary adjacency matrix in Step 4, we also built the weighted co-accessibility enhancer networks and evaluated the performance of the complexity of weighted network connectivity in predicting cell identity and disease genes. It resulted in not obvious difference between two methods (**Figure S1E**-**S1G**).

#### Quantifying network connectivity in Step 5

In **Figure S7B** and **S7C**, we quantified the network connectivity by an alternative method using the maximum degrees of nodes in network, termed the network connectivity (maximum) hereafter. Algorithmically, these two metrics, network connectivity (average) and network connectivity (maximum), are distinguished by in without or with considering the connections between non-hub enhancers.

#### Thresholds to classify enhancer networks in Step 6

To test the robustness of thresholds of network size and network connectivity in defining Complex, Multiple and Simple mode, we set different thresholds and calculated the enrichment of cell identity and disease genes (**Figure S2**).

#### The relationship of the network connectivity and network size

To decouple the network size and network connectivity, we ranked the enhancer networks based on the network size and separated them into 5 groups from high to low, which resulted in similar network size level within each group (**Figure S3A**).

Then we compared the network connectivity and cell identity/disease genes enrichment of the Complex and Multiple networks in each group (**Figure S3B-S3D**). ***The relationship of the network connectivity and chromatin accessibility***

To decouple the chromatin accessibility and network connectivity, we grouped the enhancer networks into 5 groups based on the average chromatin accessibility of the enhancers within each network from high to low, which resulted in similar chromatin accessibility level within each group (**Figure S4A**). Then we compared the network connectivity and cell identity/disease genes enrichment of the Complex and Multiple networks in each group (**Figure S4B-S4D**).

### Retrieval of cell identity and disease genes

The blood-related cell identity genes were retrieved from the website (https://www.biolegend.com/cell_markers) and (Ranzoni et al., 2021). The blood-related disease genes were from DisGeNET (Pinero et al., 2017). The skin-related cell identity genes were from (Ma et al., 2020). The skin-related disease genes were from MalaCards (https://www.malacards.org), OMIM (https://omim.org) and DisGeNET (Pinero et al., 2017). The neuron-related cell identity genes were retrieved from (Chen et al., 2019; Zhu et al., 2019). The neuron-related disease genes were from DisGeNET (Pinero et al., 2017). The kidney-related and heart-related cell identity genes were retrieved from (Domcke et al., 2020). The kidney-related and heart-related disease genes were from DisGeNET (Pinero et al., 2017). The skin-related cell identity genes were from (Ma et al., 2020). All these cell identity and disease genes are provided in Table S3.

### Enrichment analysis of cell identity and disease genes

We performed cell identity and disease genes enrichment analysis for gene groups in Complex, Multiple and Simple modes. Briefly, given a gene group, the enrichment score was calculated as the fold enrichment relative to the genome background. The computing method was determined as:

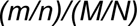

where *m* and *M* represent the number of cell identity genes within the group and genome-wide, respectively, and *n* and *N* represent the number of genes within the group and genome-wide, respectively.

### Performance evaluation in predicting cell identity and disease genes

To evaluate the performance of enhancer networks in predicting the cell identity and disease genes, we ranked all genes by various scoring methods, including network connectivity, network size, and overall chromatin accessibility. We then calculated the fold-enrichment of cell identity or disease genes in top ranked genes with a moving window of 50, using the whole genome as the background. The *p*-value, indicating the significance of the difference in performance between the two scoring methods, was determined based on the enrichment in the top 50 genes.

### Enrichment analysis of GWAS SNPs

The SNPs curated in the GWAS Catalog (Welter et al., 2014) were downloaded through the UCSC Table Browser (http://genome.ucsc.edu/). In addition, we curated a list of cell type related GWAS SNPs using a semi-automatic text mining method as described below.

#### Blood-related GWAS SNPs

The subset of blood-related GWAS SNPs was selected as those associated with at least one of the following keywords in the ‘trait’ field: ‘Erythrocyte’, ‘F-cell’, ‘HbA2’, ‘Hematocrit’, ‘Hematological’, ‘Hematology’, ‘Hemoglobin’, ‘Platelet’, ‘Blood’, ‘Anemia’, ‘Sickle cell disease’, ‘Thalassemia’, ‘Leukemia’, ‘Lymphoma’, ‘Lymphocyte’, ‘B cell ‘,‘B-cell’, ‘Lymphoma’, ‘Lymphocyte’, and ‘White blood cell’.

#### B cell-related GWAS SNPs

The subset of blood-related GWAS SNPs was selected as those associated with at least one of the following keywords in the ‘trait’ field: ‘Blood’, ‘B cell ‘, ‘B-cell’, ‘Lymphoma’, ‘Lymphocyte’.

#### Skin-related GWAS SNPs

The subset of skin-related GWAS SNPs was selected as those associated with at least one of the following keywords in the ‘trait’ field: ‘Skin’, ‘Acne’, ‘Areata’, ‘Dermatitis’, ‘Pemphigus’, ‘Psoriasis’, ‘Rosacea’, ‘Scleroderma’, ‘Vitiligo’.

#### Cerebral-related GWAS SNPs

The subset of neuron-related GWAS SNPs was selected as those associated with at least one of the following keywords in the ‘trait’ field: ‘Amyotrophic lateral sclerosis’, ‘Parkinson’s disease’, ‘Attention deficit’, ‘Anorexia’, ‘Type 1 diabetes’, ‘Ulcerative colitis’, ‘Menarche’, ‘Depressed affect’, ‘Intelligence’, ‘sclerosis’, ‘Insomnia’, ‘Menopause’, ‘Artery disease’, ‘Educational attainment’, ‘Cerebral’, ‘Ischemic’, ‘Spastic Diplegia’, ‘Malaria’, ‘Aneurysm’, ‘Cortex’, ‘Spastic Quadriplegia’, ‘Band Heterotopia’, ‘Cerebrovascular Disease’, ‘Arteriovenous Malformations of the Brain’, ‘Spastic Hemiplegia’, ‘Intracranial Embolism’, ‘Brain Edema’, ‘Brain Injury’, ‘Adrenoleukodystrophy’, ‘Intracranial Thrombosis’, ‘Seizure Disorder’, ‘Depression’, ‘Encephalopathy’, ‘Arteriovenous Malformation’, ‘Cardiac Arrest’, ‘Cerebritis’, ‘Mitochondrial DNA Depletion Syndrome 4a’, ‘Hypoxia’, ‘Thrombosis’, ‘Developmental and Epileptic Encephalopathy 39’, ‘Hemorrhage’, ‘Intracerebral’, ‘Schizophrenia’, and ‘Spasticity’.

#### Kidney-related GWAS SNPs

The subset of kidney-related GWAS SNPs was selected as those associated with at least one of the following keywords in the ‘trait’ field: ‘Kidney’, ‘Kidney Disease’, ‘nephridium’, ‘Renal’, ‘Renal Cell Carcinoma’, ‘Nonpapillary’, ‘Kidney Cancer’, ‘Autosomal Dominant Polycystic Kidney Disease’, ‘Tukel Syndrome’, ‘Leiomyosarcoma’, ‘Muscle Cancer’, ‘Smooth Muscle Tumor’, ‘Nephrolithiasis’, ‘Kidney stones’, ‘Membranous nephropathy’, ‘Urinary metabolite levels in chronic kidney disease’, ‘Estimated glomerular filtration rate’.

#### Heart-related GWAS SNPs

The subset of heart-related GWAS SNPs was selected as those associated with at least one of the following keywords in the ‘trait’ field: ‘heart Disease’, ‘Dry heart Syndrome Cataract’, ‘Fish-heart Disease’, ‘Aland Island heart Disease Cat heart Syndrome’, ‘Muscle heart Brain Disease’, ‘Ocular Cancer Myopia’, ‘Myopia’, ‘Keratoconjunctivitis Sicca’, ‘Conjunctivitis’, ‘Sjogren Syndrome’, ‘Retinal Detachment’, ‘Microvascular Complications of Diabetes 5’, ‘Open-Angle Glaucoma Refractive Error’.

#### Enrichment analysis

For each dataset, the enhancers were converted to hg38 genomic coordinates using the liftOver software from the UCSC Genome Browser (http://genome.ucsc.edu/cgi-bin/hgLiftOver). The overlap between loci and GWAS SNPs was performed using bedtools intersect (Quinlan and Hall, 2010). In short, for enhancers in each group, the enrichment score was calculated as the fold enrichment relative to the genome background. The computing method was listed as following:

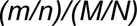

where *m* and *M* represent the number of SNPs within the group and genome-wide, respectively, and *n* and *N* represent the number of loci within the group and genome-wide, respectively. The genome-wide background is generated from a list of loci obtained by randomly shuffling the list of regular enhancers.

### Sequence conservation score

PhastCons 100-way vertebrate conservation scores were downloaded from the UCSC Genome Browser (Siepel et al., 2005). We calculated the mean PhastCons score for each enhancer as previously described (Sarropoulos et al., 2021).

### Comparison of PEIs and Hi-C chromatin interactions

High-resolution Hi-C data in GM12878 cell was obtained from the literature (Rao et al., 2014). The statistically significant chromatin interactions were detected as previously described (Huang et al., 2018). We compared the enrichment of chromatin interactions detected by Hi-C in enhancer pairs with different co-accessibility (**Figure 4A**).

### Comparison of enhancer networks based on PEIs and Hi-C chromatin interactions in GM12878 cell line

We mapped Hi-C chromatin interactions to the enhancer clusters defined by single cell GM12878 data to replace the PEIs by using bedtools map, then built enhancer networks, evaluated the complexity of enhancer networks and defined network hub enhancers following the workflow in eNet analysis.

### Trajectory analysis

We performed trajectory analysis for B cell differentiation using the method previously described (Satpathy et al., 2019) to order cells in pseudotime.

### Cell type-specific enhancer networks

To build cell type specific enhancer networks (**Figure 5**), we used the enhancer accessibility and gene expression matrix from a specific cell type as the input for eNet algorithm. The gene expression and chromatin accessibility of cell type-specific enhancer network, were represented by their average across all cells per cell type, followed by min-max normalization.

### Pseudotime difference between gene expression and enhancer networks

To compare the dynamics of gene expression enhancer networks, we quantified the difference of the pseudotime of B cell differentiation between the onset of gene expression and establishment of the enhancer network. We focused on genes highly expressed in preB or B cells and controlled by enhancer networks in Complex mode across B cell differentiation pseudotime. First, for each single cell, we assigned the gene expression, network connectivity, and chromatin accessibility based on their cell type annotations, which were further smoothed by applying a sliding window of 50 cells along the pseudotime. We then defined the time of gene expression onset and enhancer network establishment, measured by chromatin accessibility or network connections, at the first instance of the smoothed value being larger than the predefined value. Finally, the pseudotime lag was calculated as the time of gene expression onset subtracted by the time of enhancer network establishment.

### Blood-related SEs

The SEs list associated with blood-related cell types from the dbSUPER database (Khan and Zhang, 2016) was curated into a catalog of blood-related SEs (**Table S4**). We first downloaded the corresponding SE list from dbSUPER, sorted, and merged into an SE list using bedtools (Quinlan and Hall, 2010). In this way, we generated 2,306 human blood-related SEs in total.

### Gene Ontology (GO) enrichment analysis

Gene Ontology (GO) enrichment analysis of enhancer network target genes was performed by clusterProfiler package (Yu et al., 2012).

## Data availability

All datasets analyzed in this study were published previously. The corresponding descriptions and GEO number are described in the **Table S1**.

## Code availability

The full code of eNet was provided in the **Supplementary material** and made available via GitHub, see https://github.com/xmuhuanglab/eNet.

## Authors’ contributions

D.H. and J.H. conceived and designed the study. D.H., H.L., L.L. and M.T. performed the computational analysis. D.H., H.L., L.L., M.S., J.D., F.L. and J.H. wrote the manuscript. J.H. supervised the study.

## Acknowledgements

We thank the useful comments from Drs. Guo-Cheng Yuan, Jian Xu and the members in J.H. lab. This work was supported by the National Natural Science Foundation of China (31871317, 32070635, 82002529, 81891000 and 81891002) and Natural Science Foundation of Fujian Province of China (2020J01028 and 2020J05012), and the Strategic Priority Research Program of the Chinese Academy of Sciences (XDA16040000).

## Supplemental Figures and Legends

**Figure S1.**
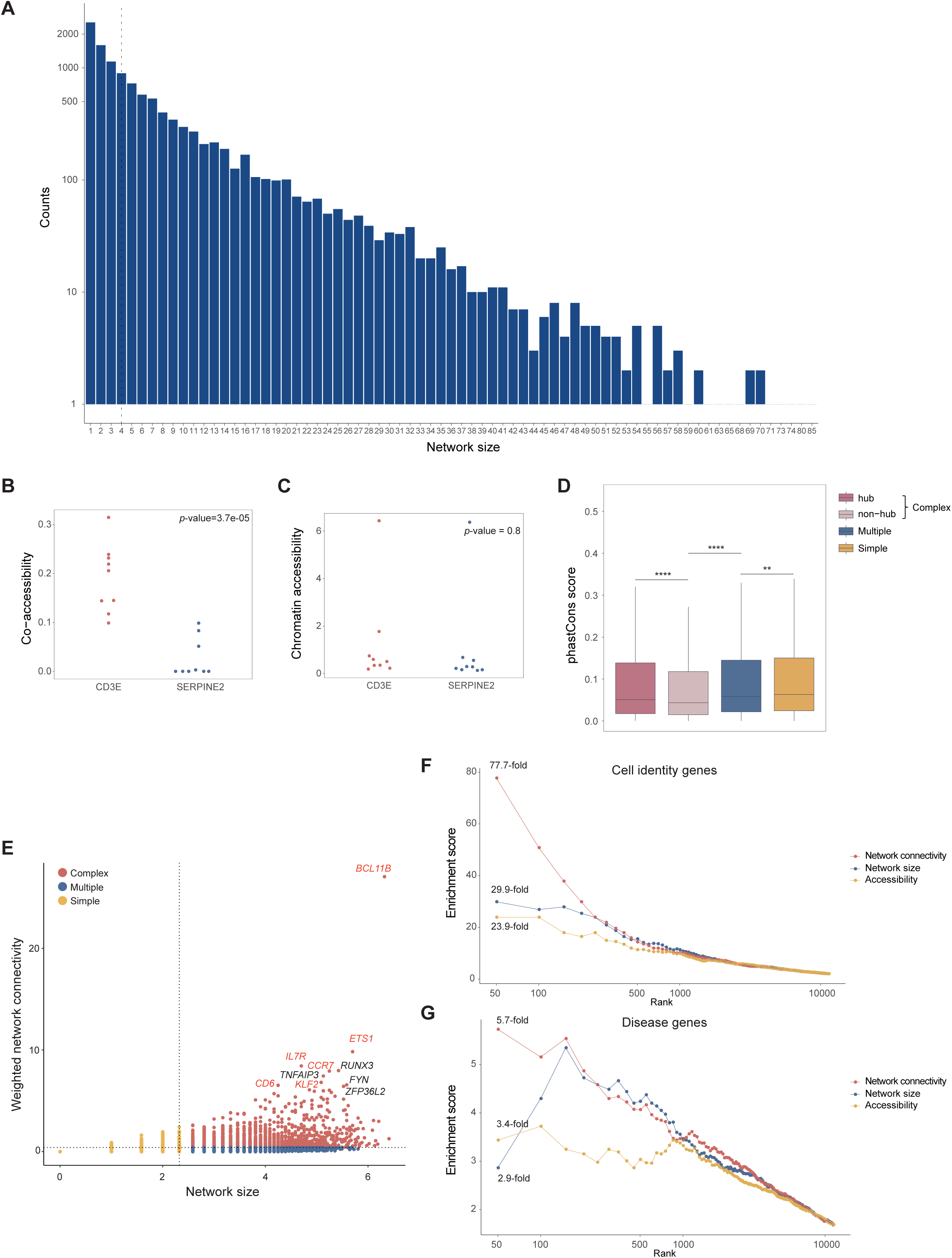
Enhancer networks during human hematopoiesis, related to Figure 2. **(A)** Distribution of network size for the enhancer networks during human hematopoiesis, where the dash line indicates the median of network size. **(B** and **C)** The co-accessibility (**B**) and chromatin accessibility (**C**) of the constituent enhancers in the example of Fig 2C. *p-*values were calculated using Student’s *t*-test. (D) PhastCons conservation score of enhancers in Complex (hub and non-hub), Multiple and Simple groups. (E) Scatter plot of the weighted enhancer networks during hematopoiesis, where the x-axis represents the network size (log2-scaled) and the y-axis represents network connectivity. Top 10 genes ranked by network connectivity were labelled, where known blood-related cell identity or disease genes were red-highlighted. (**F** and **G**) Enrichment of cell identity (**F**) and disease genes (**G**) (y-axis) is plotted for top genes (x-axis) ranked by different properties of the weighted enhancer networks in (**E**), including network connectivity (the frequency of PEIs in this study), network size (equivalent to the enhancer number in multiple enhancers, or overall chromatin accessibility of enhancers.

**Figure S2.**
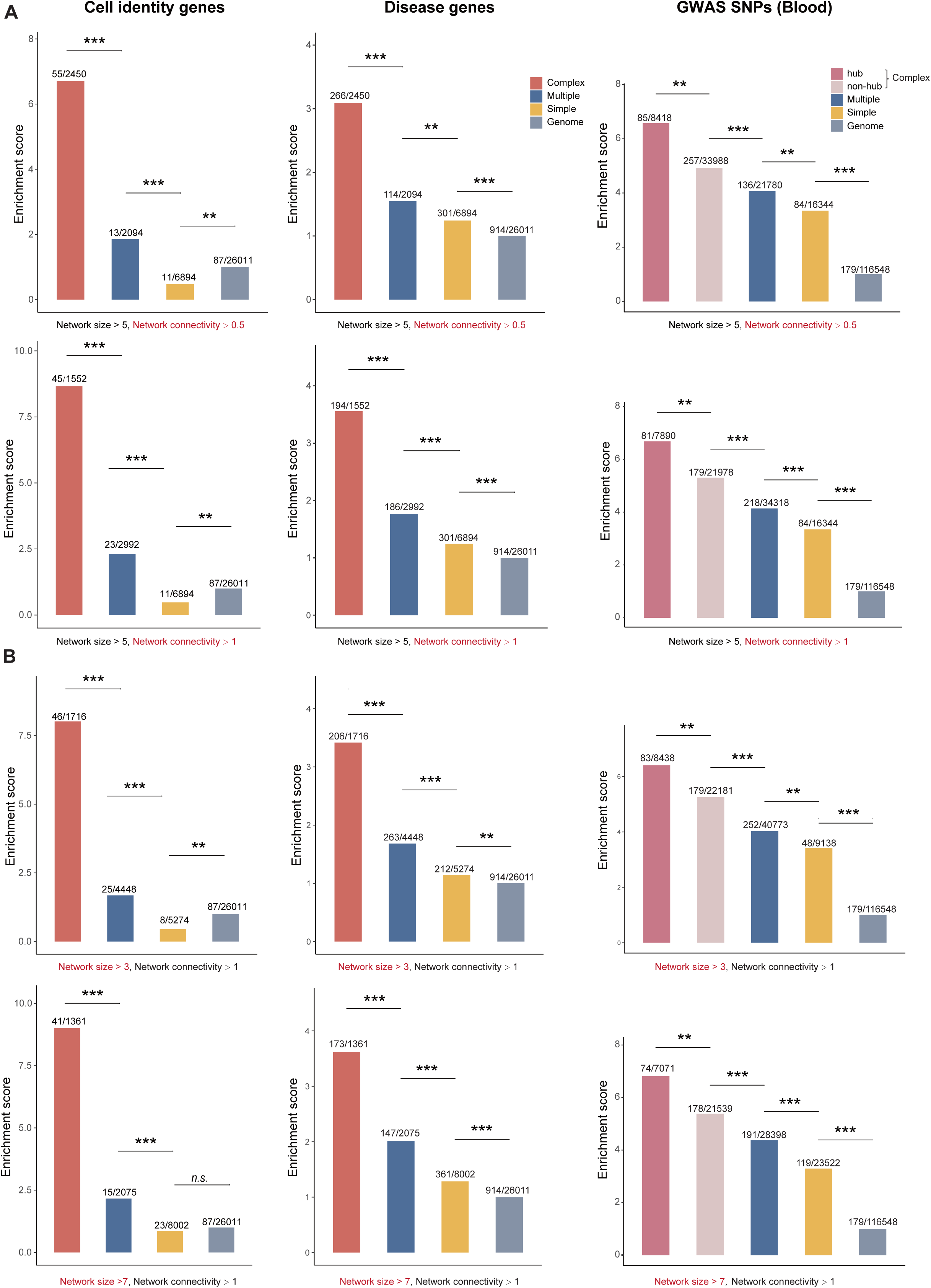
Classification of the enhancer networks using various threshold values of network connectivity and network size in human blood dataset, related to Figure 2. (**A** and **B)** Various threshold values of network connectivity (**A**) and network size (**B**). Enrichment of cell identity (left) and disease genes (middle) in genes in Complex, Multiple, and Simple modes, using the whole genome as the background. Enrichment of the diseases/traits-related SNPs curated in the GWAS catalog (right) in enhancers in Complex (hub and non-hub), Multiple, and Simple modes, using randomly selected genomic regions as the control. *p*-values were calculated using the binomial test. \**p* < 0.05; \*\**p* < 0.01; \*\*\**p* < 0.001; *n.s.,* not significant.

**Figure S3.**
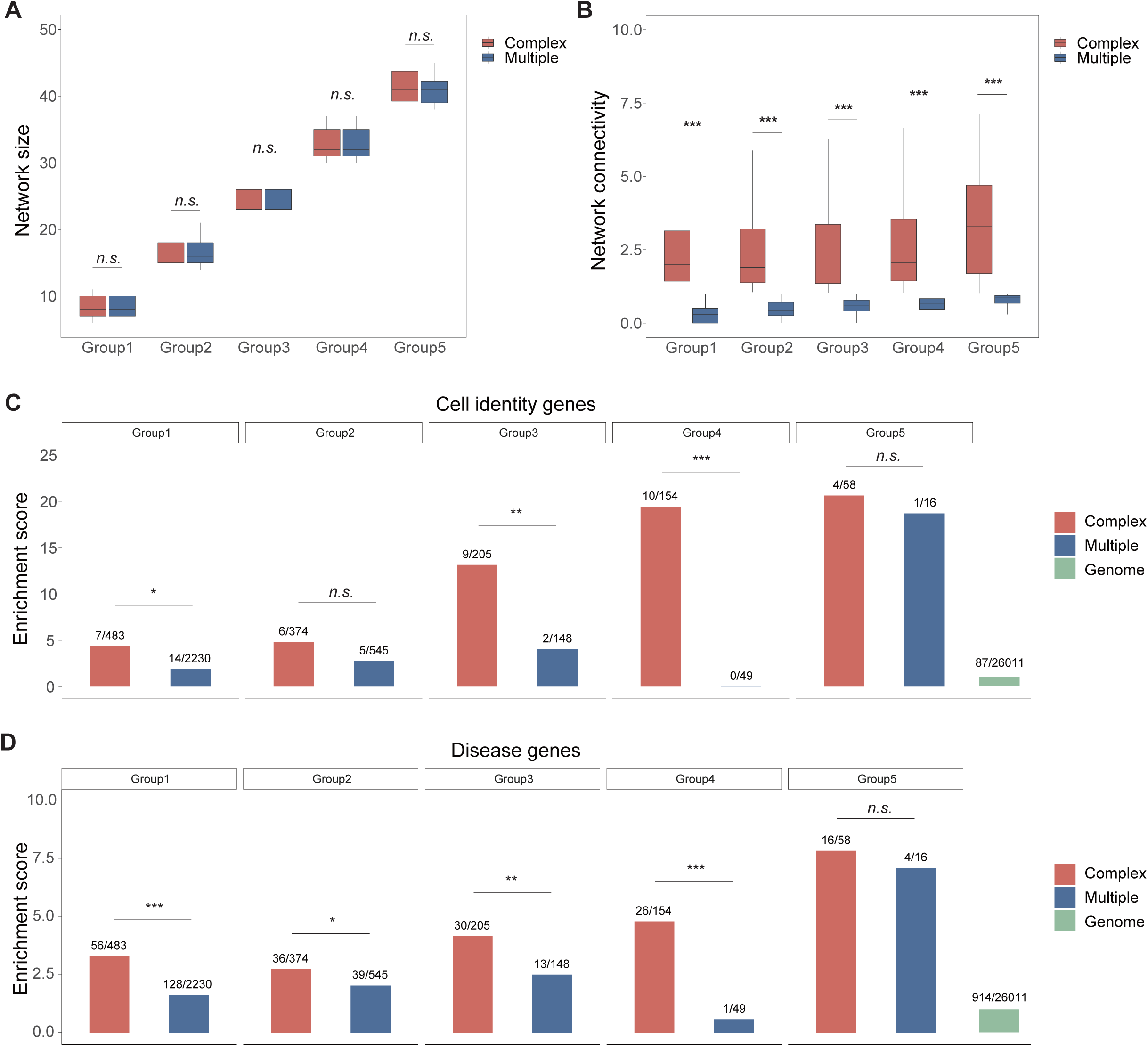
The relationship of the network connectivity and network size, related to Figure 2. (**A** and **B)** The network size (**A**) and network connectivity (**B**) in genes regulated by Complex and Multiple enhancer networks with a similar network size level. *p-*values were calculated using Student’s *t*-test. \**p* < 0.05; \*\**p* < 0.01; \*\*\**p* < 0.001; *n.s.,* not significant. **(C** and **D)** Enrichment of blood-related cell identity genes (**C**) and disease genes (**D**) in genes regulated by Complex and Multiple enhancer networks with a similar network size level, using whole genome as the background. *p*-values were calculated using the binomial test. \**p* < 0.05; \*\**p* < 0.01; \*\*\**p* < 0.001; *n.s.,* not significant.

**Figure S4.**
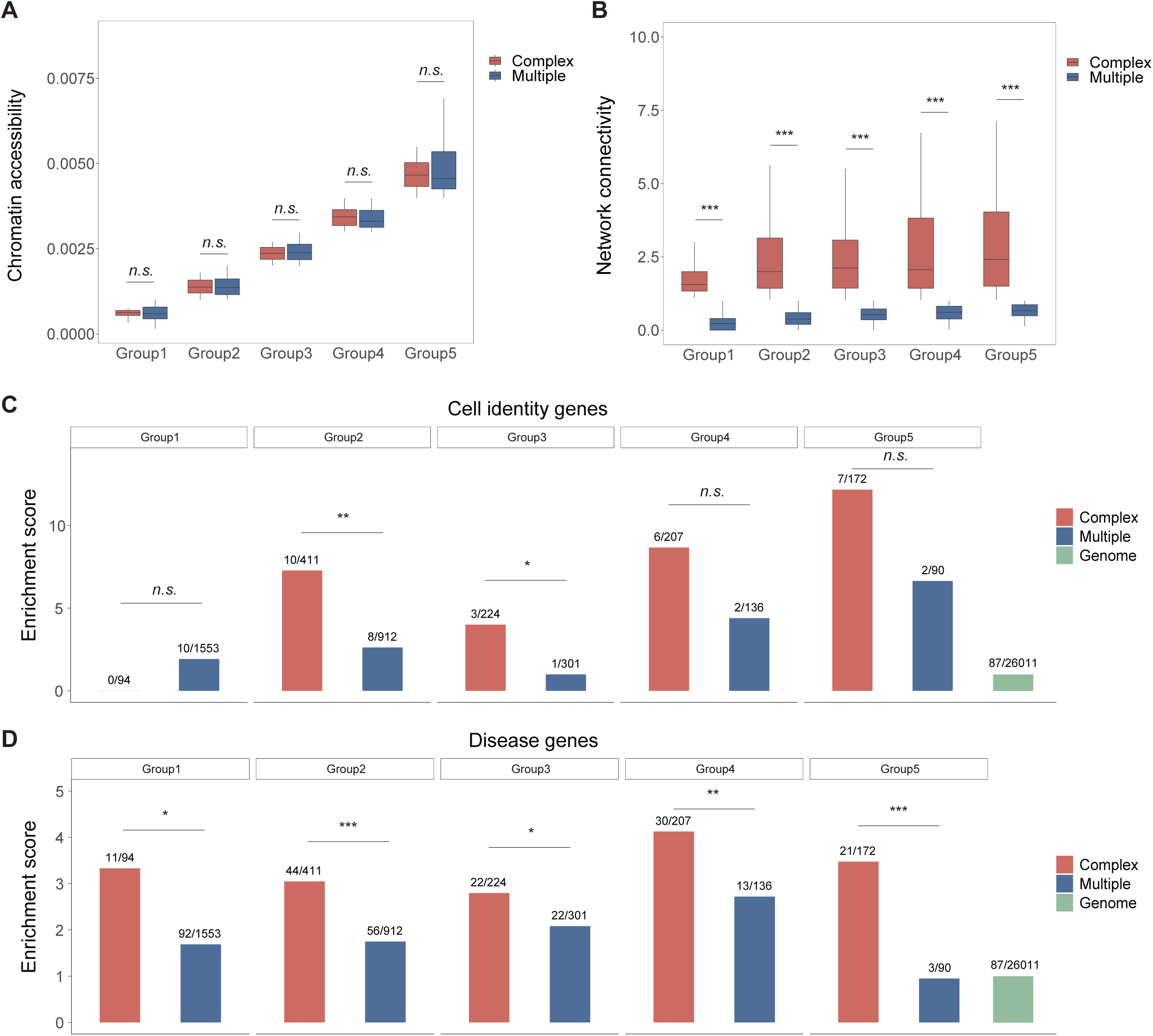
The relationship of the network connectivity and chromatin accessibility, related to Figure 2. (**A** and **B)** The chromatin accessibility (**A**) and network connectivity (**B**) in genes regulated by Complex and Multiple enhancer networks with a similar chromatin accessibility level. *p-*values were calculated using Student’s *t*-test. \**p* < 0.05; \*\**p* < 0.01; \*\*\**p* < 0.001; *n.s.,* not significant. **(C** and **D)** Enrichment of blood-related cell identity genes (**C**) and disease genes (**D**) in genes regulated by Complex and Multiple enhancer networks with a similar chromatin accessibility level, using whole genome as the background. *p*-values were calculated using the binomial test. \**p* < 0.05; \*\**p* < 0.01; \*\*\**p* < 0.001; *n.s.,* not significant.

**Figure S5.**
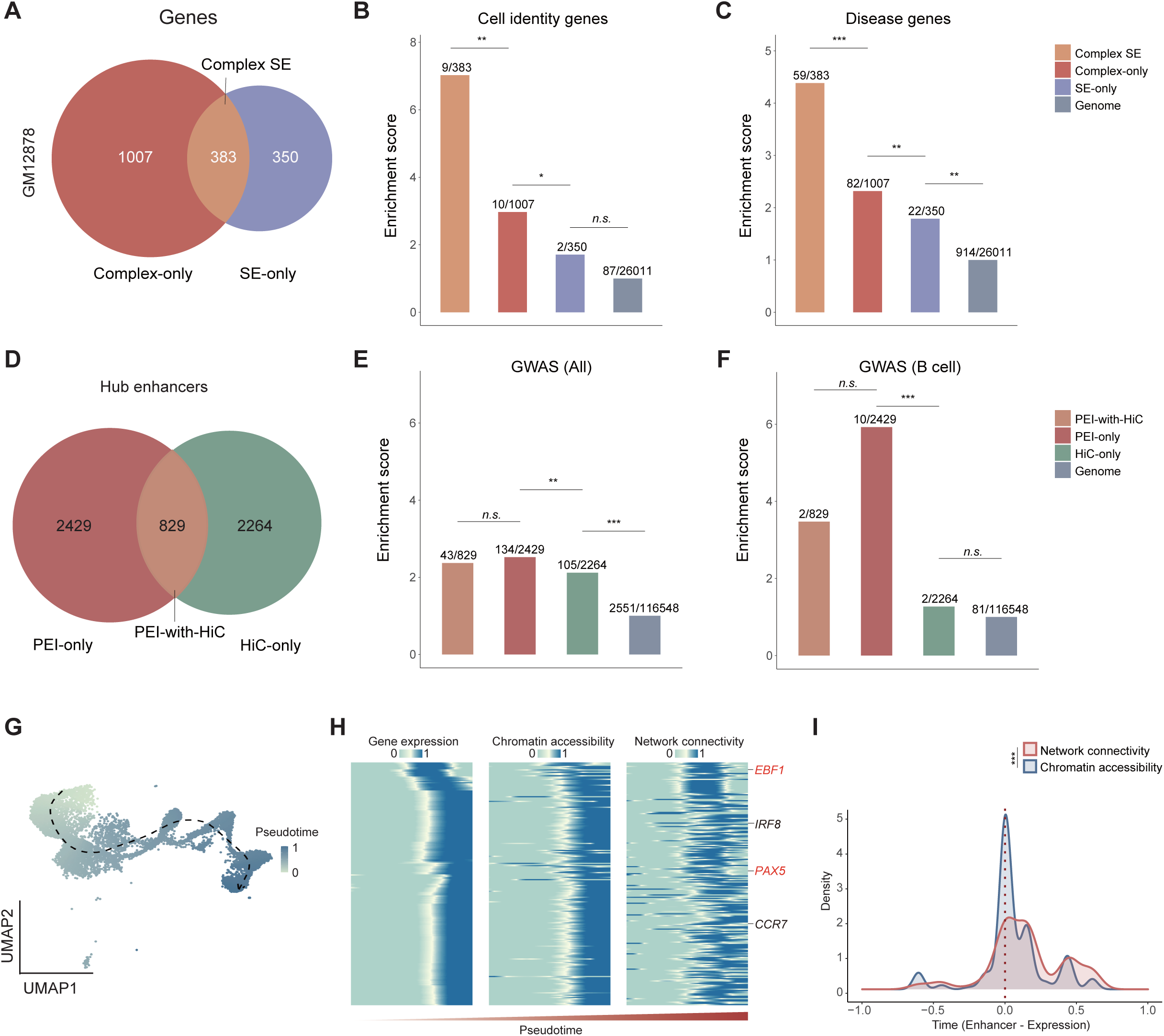
The SEs and Hi-C in GM12878 cell line and dynamic of enhancer networks during B cell differentiation, related to Figure 3-5. **(A)** Venn diagram showing the overlap of genes regulated by enhancer networks in Complex mode and SEs in GM12878 dataset, resulting in three groups: Complex SE, Complex-only, and SE-only. **(B** and **C)** Enrichment of cell identity (**B**) and disease genes (**C**) in genes in three groups, using the whole genome as the background. *p-*values were calculated using the binomial test. *\*p* < 0.05*; **p* < 0.01*; ***p* < 0.001; *n.s.,* not significant. (**D**) Venn diagram showing the overlap between the network hub enhancers based on PEI and Hi-C data in the enhancer networks in PEI-with-HiC group in Figure 4D. (**E** and **F**) Enrichment of all GWAS SNPs (**E**) and B cell-related GWAS SNPs (**F**) in three groups of hub enhancers: PEI-only, Hi-C-only and PEI-with-Hi-C. *p*-values were calculated using the binomial test. \**p* < 0.05; \*\**p* < 0.01; \*\*\**p* < 0.001; *n.s.*, not significant. **(G)** The pseudotime during B cell differentiation inferred based on scATAC-seq data. **(H)** Dynamics of gene expression (left), chromatin accessibility (middle), and enhancer network connectivity (right) across the B cell differentiation pseudotime (column). Genes highly expressed in pre-B or B cells and controlled by enhancer networks in the Complex mode are included (row), where some known cell identity genes are labelled. **(I)** Difference of the pseudotime of B cell differentiation between onset of gene expression and establishment of enhancer networks. *p-*values were calculated using one-sided paired Student’s *t*-test. *\*p* < 0.05*; **p* < 0.01*; ***p* < 0.001; *n.s.,* not significant.

**Figure S6.**
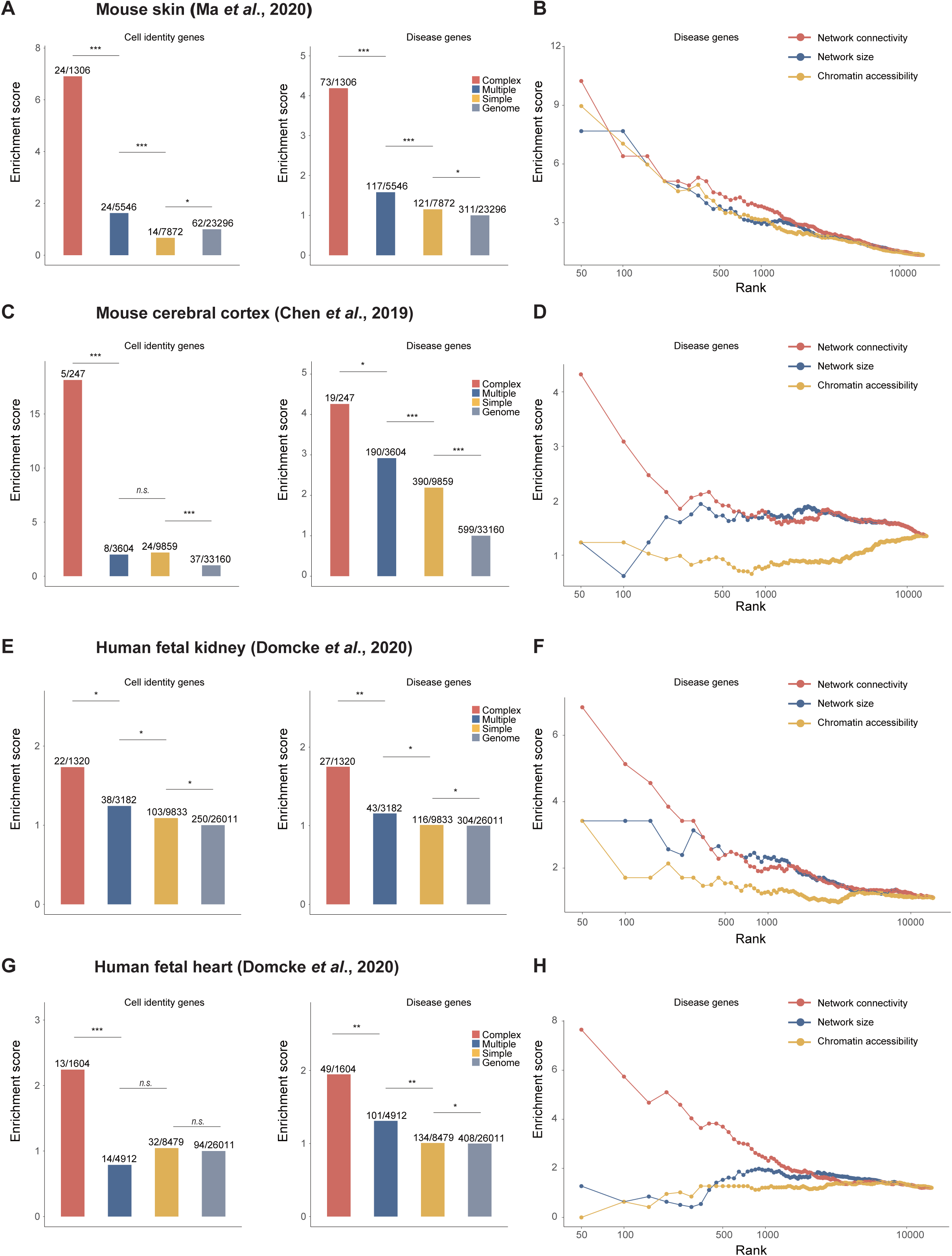
eNet analysis in various human or mouse tissues across different single-cell platforms, related to Figure 6. **(A, C, E, G)** Enrichment of cell identity (left) and disease genes (right) in genes in Complex, Multiple and Simple modes, using the whole genome as the background. (**A**) mouse skin dataset (SHARE-seq) (Ma et al., 2020), (**C**) mouse cerebral cortex dataset (SNARE-seq) (Chen et al., 2019), (**E**) human fetal kidney dataset (sci-ATAC-seq3) (Domcke et al., 2020), and (**G**) human fetal heart dataset (sci-ATAC-seq3) (Domcke et al., 2020). The number of cell identity or disease genes and total genes in each group are labelled on each bar. p-values were calculated using the binomial test. \**p* < 0.05; \*\**p* < 0.01; \*\*\**p* < 0.001; n.s., not significant. **(B, D, F, H)** Enrichment of disease genes (y-axis) is plotted for top genes ranked by various scoring methods (x-axis) in different tissues and approaches. (**B**) mouse skin dataset, (**D**) mouse cerebral cortex dataset, (**F**) human fetal kidney dataset, and (**H**) human fetal heart dataset.

**Figure S7.**
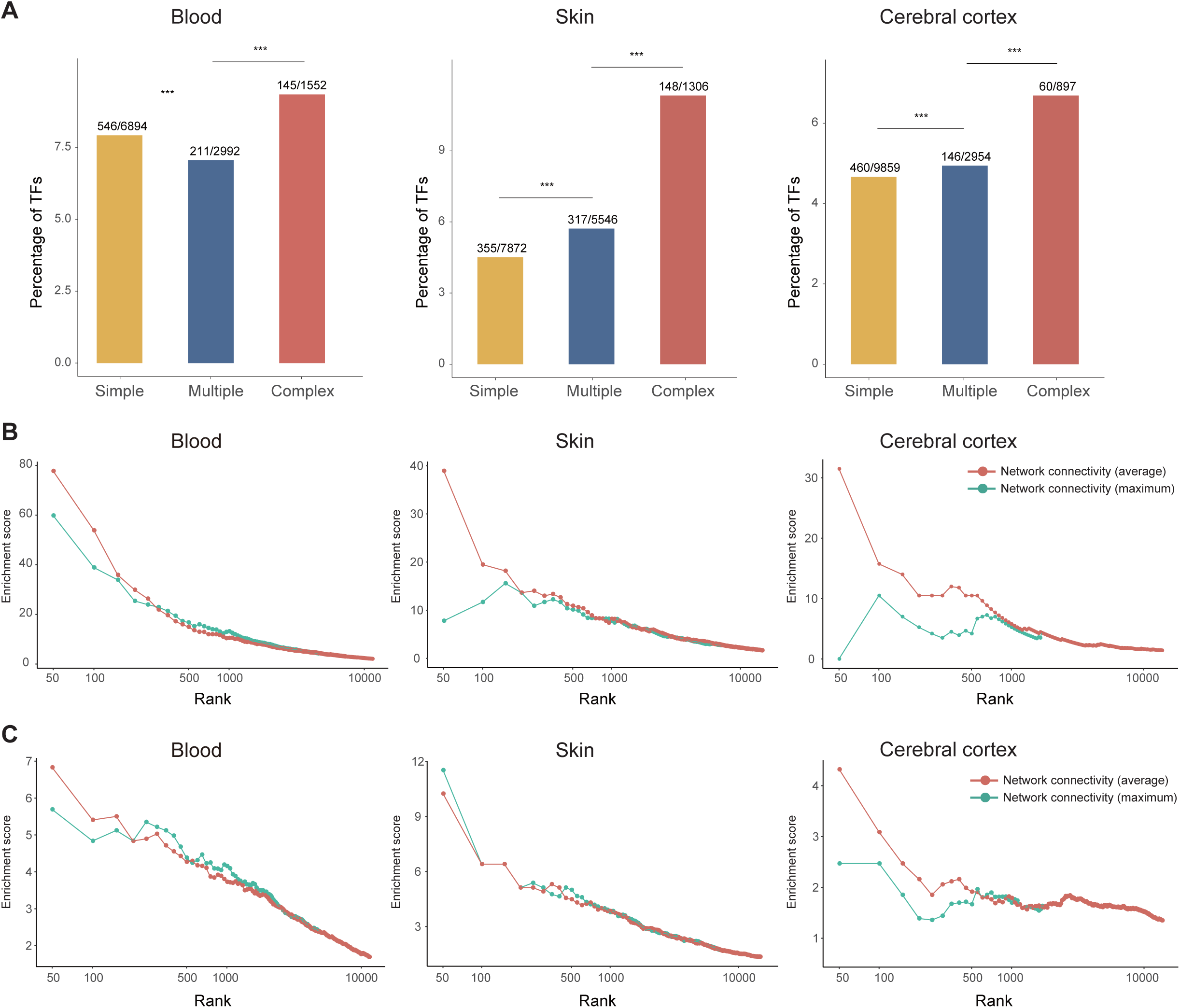
Three modes of enhancer networks in gene regulation, related to Figure 7. **(A)** Percentage of transcription factors (TFs) in genes regulated by enhancer networks in the Simple, Multiple and Complex modes. *p-*values were calculated using the binomial-test. *\*p* < 0.05*; **p* < 0.01*; ***p* < 0.001; *n.s.,* not significant. **(B** and **C)** Enrichment of cell identity (**B**) and disease genes (**C**) ranked by two network connectivity metrics: enhancer network connectivity (average) and network connectivity (maximum) in the human blood, mouse skin, and mouse cerebral cortex datasets.

## Supplementary Tables

**Table S1**

A list of all used datasets and accession numbers, related to Figure 2-6.

**Table S2**

List of enhancer networks in human blood, mouse skin, cerebral cortex, human fetal kidney and heart datasets, related to Figure 2-6.

**Table S3**

The list of cell identity and disease genes for human blood, mouse skin, cerebral cortex, human fetal kidney and heart datasets, related to Figure 2-6.

**Table S4**

The list of blood-related SEs catalog, related to Figure 3 and 4.

## References

Argelaguet, R., Clark, S.J., Mohammed, H., Stapel, L.C., Krueger, C., Kapourani, C.A., Imaz-Rosshandler, I., Lohoff, T., Xiang, Y., Hanna, C.W., et al. (2019). Multi-omics profiling of mouse gastrulation at single-cell resolution. Nature 576, 487–491.

Bahr, C., von Paleske, L., Uslu, V.V., Remeseiro, S., Takayama, N., Ng, S.W., Murison, A., Langenfeld, K., Petretich, M., Scognamiglio, R., et al. (2018). A Myc enhancer cluster regulates normal and leukaemic haematopoietic stem cell hierarchies. Nature 553, 515–520.

Banerji, J., Rusconi, S., and Schaffner, W. (1981). Expression of a beta-globin gene is enhanced by remote SV40 DNA sequences. Cell 27, 299–308.

Barabasi, A.-L. (2016). Network Science (Cambridge University Press).

Blobel, G.A., Higgs, D.R., Mitchell, J.A., Notani, D., and Young, R.A. (2021). Testing the super-enhancer concept. Nat Rev Genet.

Buenrostro, J.D., Wu, B., Litzenburger, U.M., Ruff, D., Gonzales, M.L., Snyder, M.P., Chang, H.Y., and Greenleaf, W.J. (2015). Single-cell chromatin accessibility reveals principles of regulatory variation. Nature 523, 486–490.

Cai, W., Huang, J., Zhu, Q., Li, B.E., Seruggia, D., Zhou, P., Nguyen, M., Fujiwara, Y., Xie, H., Yang, Z., et al. (2020). Enhancer dependence of cell-type-specific gene expression increases with developmental age. Proc Natl Acad Sci U S A 117, 21450–21458.

Canver, M.C., Smith, E.C., Sher, F., Pinello, L., Sanjana, N.E., Shalem, O., Chen, D.D., Schupp, P.G., Vinjamur, D.S., Garcia, S.P., et al. (2015). BCL11A enhancer dissection by Cas9-mediated in situ saturating mutagenesis. Nature 527, 192–197.

Cao, J., Cusanovich, D.A., Ramani, V., Aghamirzaie, D., Pliner, H.A., Hill, A.J., Daza, R.M., McFaline-Figueroa, J.L., Packer, J.S., Christiansen, L., et al. (2018). Joint profiling of chromatin accessibility and gene expression in thousands of single cells. Science 361, 1380–1385.

Cao, J., O’Day, D.R., Pliner, H.A., Kingsley, P.D., Deng, M., Daza, R.M., Zager, M.A., Aldinger, K.A., Blecher-Gonen, R., Zhang, F., et al. (2020). A human cell atlas of fetal gene expression. Science 370.

Carleton, J.B., Berrett, K.C., and Gertz, J. (2017). Multiplex Enhancer Interference Reveals Collaborative Control of Gene Regulation by Estrogen Receptor alpha-Bound Enhancers. Cell Syst 5, 333–344 e335.

Chen, S., Lake, B.B., and Zhang, K. (2019). High-throughput sequencing of the transcriptome and chromatin accessibility in the same cell. Nat Biotechnol 37, 1452–1457.

Chen, X., Zhou, J., Zhang, R., Wong, A.K., Park, C.Y., Theesfeld, C.L., and Troyanskaya, O.G. (2021). Tissue-specific enhancer functional networks for associating distal regulatory regions to disease. Cell Syst 12, 353–362 e356.

Consortium, E.P. (2012). An integrated encyclopedia of DNA elements in the human genome. Nature 489, 57–74.

Dixon, J.R., Selvaraj, S., Yue, F., Kim, A., Li, Y., Shen, Y., Hu, M., Liu, J.S., and Ren, B. (2012). Topological domains in mammalian genomes identified by analysis of chromatin interactions. Nature 485, 376–380.

Domcke, S., Hill, A.J., Daza, R.M., Cao, J., O’Day, D.R., Pliner, H.A., Aldinger, K.A., Pokholok, D., Zhang, F., Milbank, J.H., et al. (2020). A human cell atlas of fetal chromatin accessibility. Science 370.

Fulco, C.P., Munschauer, M., Anyoha, R., Munson, G., Grossman, S.R., Perez, E.M., Kane, M., Cleary, B., Lander, E.S., and Engreitz, J.M. (2016). Systematic mapping of functional enhancer-promoter connections with CRISPR interference. Science 354, 769–773.

Gasperini, M., Tome, J.M., and Shendure, J. (2020). Towards a comprehensive catalogue of validated and target-linked human enhancers. Nat Rev Genet.

Gonzalez, A.J., Setty, M., and Leslie, C.S. (2015). Early enhancer establishment and regulatory locus complexity shape transcriptional programs in hematopoietic differentiation. Nat Genet 47, 1249–1259.

Granja, J.M., Klemm, S., McGinnis, L.M., Kathiria, A.S., Mezger, A., Corces, M.R., Parks, B., Gars, E., Liedtke, M., Zheng, G.X.Y., et al. (2019). Single-cell multiomic analysis identifies regulatory programs in mixed-phenotype acute leukemia. Nat Biotechnol 37, 1458–1465.

Hay, D., Hughes, J.R., Babbs, C., Davies, J.O.J., Graham, B.J., Hanssen, L., Kassouf, M.T., Marieke Oudelaar, A.M., Sharpe, J.A., Suciu, M.C., et al. (2016). Genetic dissection of the alpha-globin super-enhancer in vivo. Nat Genet 48, 895–903.

Hnisz, D., Abraham, B.J., Lee, T.I., Lau, A., Saint-Andre, V., Sigova, A.A., Hoke, H.A., and Young, R.A. (2013). Super-enhancers in the control of cell identity and disease. Cell 155, 934–947.

Huang, J., Li, K., Cai, W., Liu, X., Zhang, Y., Orkin, S.H., Xu, J., and Yuan, G.C. (2018). Dissecting super-enhancer hierarchy based on chromatin interactions. Nat Commun 9, 943.

Huang, J., Liu, X., Li, D., Shao, Z., Cao, H., Zhang, Y., Trompouki, E., Bowman, T.V., Zon, L.I., Yuan, G.C., et al. (2016). Dynamic Control of Enhancer Repertoires Drives Lineage and Stage-Specific Transcription during Hematopoiesis. Dev Cell 36, 9–23.

Huang, J., Marco, E., Pinello, L., and Yuan, G.C. (2015). Predicting chromatin organization using histone marks. Genome Biol 16, 162.

Jinek, M., Chylinski, K., Fonfara, I., Hauer, M., Doudna, J.A., and Charpentier, E. (2012). A programmable dual-RNA-guided DNA endonuclease in adaptive bacterial immunity. Science 337, 816–821.

Kai, Y., Li, B.E., Zhu, M., Li, G.Y., Chen, F., Han, Y., Cha, H.J., Orkin, S.H., Cai, W., Huang, J., et al. (2021). Mapping the evolving landscape of super-enhancers during cell differentiation. Genome Biol 22, 269.

Khan, A., and Zhang, X. (2016). dbSUPER: a database of super-enhancers in mouse and human genome. Nucleic Acids Res 44, D164–171.

Lambert, S.A., Jolma, A., Campitelli, L.F., Das, P.K., Yin, Y., Albu, M., Chen, X., Taipale, J., Hughes, T.R., and Weirauch, M.T. (2018). The Human Transcription Factors. Cell 172, 650–665.

Lara-Astiaso, D., Weiner, A., Lorenzo-Vivas, E., Zaretsky, I., Jaitin, D.A., David, E., Keren-Shaul, H., Mildner, A., Winter, D., Jung, S., et al. (2014). Immunogenetics. Chromatin state dynamics during blood formation. Science 345, 943–949.

Li, Y.E., Preissl, S., Hou, X., Zhang, Z., Zhang, K., Qiu, Y., Poirion, O.B., Li, B., Chiou, J., Liu, H., et al. (2021). An atlas of gene regulatory elements in adult mouse cerebrum. Nature 598, 129–136.

Lieberman-Aiden, E., van Berkum, N.L., Williams, L., Imakaev, M., Ragoczy, T., Telling, A., Amit, I., Lajoie, B.R., Sabo, P.J., Dorschner, M.O., et al. (2009). Comprehensive mapping of long-range interactions reveals folding principles of the human genome. Science 326, 289–293.

Liu, X., Chen, Y., Zhang, Y., Liu, Y., Liu, N., Botten, G.A., Cao, H., Orkin, S.H., Zhang, M.Q., and Xu, J. (2020). Multiplexed capture of spatial configuration and temporal dynamics of locus-specific 3D chromatin by biotinylated dCas9. Genome Biol 21, 59.

Liu, X., Zhang, Y., Chen, Y., Li, M., Zhou, F., Li, K., Cao, H., Ni, M., Liu, Y., Gu, Z., et al. (2017). In Situ Capture of Chromatin Interactions by Biotinylated dCas9. Cell 170, 1028–1043 e1019.

Long, H.K., Prescott, S.L., and Wysocka, J. (2016). Ever-Changing Landscapes: Transcriptional Enhancers in Development and Evolution. Cell 167, 1170–1187.

Ma, S., Zhang, B., LaFave, L.M., Earl, A.S., Chiang, Z., Hu, Y., Ding, J., Brack, A., Kartha, V.K., Tay, T., et al. (2020). Chromatin Potential Identified by Shared Single-Cell Profiling of RNA and Chromatin. Cell 183, 1103–1116 e1120.

Malin, J., Aniba, M.R., and Hannenhalli, S. (2013). Enhancer networks revealed by correlated DNAse hypersensitivity states of enhancers. Nucleic Acids Res 41, 6828–6838.

Maurano, M.T., Humbert, R., Rynes, E., Thurman, R.E., Haugen, E., Wang, H., Reynolds, A.P., Sandstrom, R., Qu, H., Brody, J., et al. (2012). Systematic localization of common disease-associated variation in regulatory DNA. Science 337, 1190–1195.

Mumbach, M.R., Satpathy, A.T., Boyle, E.A., Dai, C., Gowen, B.G., Cho, S.W., Nguyen, M.L., Rubin, A.J., Granja, J.M., Kazane, K.R., et al. (2017). Enhancer connectome in primary human cells identifies target genes of disease-associated DNA elements. Nat Genet 49, 1602–1612.

Neph, S., Vierstra, J., Stergachis, A.B., Reynolds, A.P., Haugen, E., Vernot, B., Thurman, R.E., John, S., Sandstrom, R., Johnson, A.K., et al. (2012). An expansive human regulatory lexicon encoded in transcription factor footprints. Nature 489, 83–90.

Osterwalder, M., Barozzi, I., Tissieres, V., Fukuda-Yuzawa, Y., Mannion, B.J., Afzal, S.Y., Lee, E.A., Zhu, Y., Plajzer-Frick, I., Pickle, C.S., et al. (2018). Enhancer redundancy provides phenotypic robustness in mammalian development. Nature 554, 239–243.

Perry, M.W., Boettiger, A.N., and Levine, M. (2011). Multiple enhancers ensure precision of gap gene-expression patterns in the Drosophila embryo. Proc Natl Acad Sci U S A 108, 13570–13575.

Pinero, J., Bravo, A., Queralt-Rosinach, N., Gutierrez-Sacristan, A., Deu-Pons, J., Centeno, E., Garcia-Garcia, J., Sanz, F., and Furlong, L.I. (2017). DisGeNET: a comprehensive platform integrating information on human disease-associated genes and variants. Nucleic Acids Res 45, D833–D839.

Pliner, H.A., Packer, J.S., McFaline-Figueroa, J.L., Cusanovich, D.A., Daza, R.M., Aghamirzaie, D., Srivatsan, S., Qiu, X., Jackson, D., Minkina, A., et al. (2018). Cicero Predicts cis-Regulatory DNA Interactions from Single-Cell Chromatin Accessibility Data. Mol Cell 71, 858–871 e858.

Pott, S., and Lieb, J.D. (2015). What are super-enhancers? Nat Genet 47, 8–12.

Quinlan, A.R., and Hall, I.M. (2010). BEDTools: a flexible suite of utilities for comparing genomic features. Bioinformatics 26, 841–842.

Rada-Iglesias, A., Bajpai, R., Swigut, T., Brugmann, S.A., Flynn, R.A., and Wysocka, J. (2011). A unique chromatin signature uncovers early developmental enhancers in humans. Nature 470, 279–283.

Ranzoni, A.M., Tangherloni, A., Berest, I., Riva, S.G., Myers, B., Strzelecka, P.M., Xu, J., Panada, E., Mohorianu, I., Zaugg, J.B., et al. (2021). Integrative Single-Cell RNA-Seq and ATAC-Seq Analysis of Human Developmental Hematopoiesis. Cell Stem Cell 28, 472–487 e477.

Rao, S.S., Huntley, M.H., Durand, N.C., Stamenova, E.K., Bochkov, I.D., Robinson, J.T., Sanborn, A.L., Machol, I., Omer, A.D., Lander, E.S., et al. (2014). A 3D map of the human genome at kilobase resolution reveals principles of chromatin looping. Cell 159, 1665–1680.

Sabari, B.R., Dall’Agnese, A., Boija, A., Klein, I.A., Coffey, E.L., Shrinivas, K., Abraham, B.J., Hannett, N.M., Zamudio, A.V., Manteiga, J.C., et al. (2018). Coactivator condensation at super-enhancers links phase separation and gene control. Science 361.

Sarropoulos, I., Sepp, M., Fromel, R., Leiss, K., Trost, N., Leushkin, E., Okonechnikov, K., Joshi, P., Giere, P., Kutscher, L.M., et al. (2021). Developmental and evolutionary dynamics of cis-regulatory elements in mouse cerebellar cells. Science 373.

Satpathy, A.T., Granja, J.M., Yost, K.E., Qi, Y., Meschi, F., McDermott, G.P., Olsen, B.N., Mumbach, M.R., Pierce, S.E., Corces, M.R., et al. (2019). Massively parallel single-cell chromatin landscapes of human immune cell development and intratumoral T cell exhaustion. Nat Biotechnol 37, 925–936.

Schmitt, A.D., Hu, M., Jung, I., Xu, Z., Qiu, Y., Tan, C.L., Li, Y., Lin, S., Lin, Y., Barr, C.L., et al. (2016). A Compendium of Chromatin Contact Maps Reveals Spatially Active Regions in the Human Genome. Cell Rep 17, 2042–2059.

Schoenfelder, S., and Fraser, P. (2019). Long-range enhancer-promoter contacts in gene expression control. Nat Rev Genet 20, 437–455.

Shin, H.Y., Willi, M., HyunYoo, K., Zeng, X., Wang, C., Metser, G., and Hennighausen, L. (2016). Hierarchy within the mammary STAT5-driven Wap super-enhancer. Nat Genet 48, 904–911.

Siepel, A., Bejerano, G., Pedersen, J.S., Hinrichs, A.S., Hou, M., Rosenbloom, K., Clawson, H., Spieth, J., Hillier, L.W., Richards, S., et al. (2005). Evolutionarily conserved elements in vertebrate, insect, worm, and yeast genomes. Genome Res 15, 1034–1050.

Song, M., Pebworth, M.P., Yang, X., Abnousi, A., Fan, C., Wen, J., Rosen, J.D., Choudhary, M.N.K., Cui, X., Jones, I.R., et al. (2020). Cell-type-specific 3D epigenomes in the developing human cortex. Nature 587, 644–649.

Tang, F., Barbacioru, C., Wang, Y., Nordman, E., Lee, C., Xu, N., Wang, X., Bodeau, J., Tuch, B.B., Siddiqui, A., et al. (2009). mRNA-Seq whole-transcriptome analysis of a single cell. Nat Methods 6, 377–382.

Tang, Z., Luo, O.J., Li, X., Zheng, M., Zhu, J.J., Szalaj, P., Trzaskoma, P., Magalska, A., Wlodarczyk, J., Ruszczycki, B., et al. (2015). CTCF-Mediated Human 3D Genome Architecture Reveals Chromatin Topology for Transcription. Cell 163, 1611–1627.

Thomas, H.F., Kotova, E., Jayaram, S., Pilz, A., Romeike, M., Lackner, A., Penz, T., Bock, C., Leeb, M., Halbritter, F., et al. (2021). Temporal dissection of an enhancer cluster reveals distinct temporal and functional contributions of individual elements. Mol Cell 81, 969–982 e913.

Trevino, A.E., Muller, F., Andersen, J., Sundaram, L., Kathiria, A., Shcherbina, A., Farh, K., Chang, H.Y., Pasca, A.M., Kundaje, A., et al. (2021). Chromatin and gene-regulatory dynamics of the developing human cerebral cortex at single-cell resolution. Cell.

Tsai, A., Alves, M.R., and Crocker, J. (2019). Multi-enhancer transcriptional hubs confer phenotypic robustness. Elife 8.

Welter, D., MacArthur, J., Morales, J., Burdett, T., Hall, P., Junkins, H., Klemm, A., Flicek, P., Manolio, T., Hindorff, L., et al. (2014). The NHGRI GWAS Catalog, a curated resource of SNP-trait associations. Nucleic Acids Res 42, D1001–1006.

Yu, G., Wang, L.G., Han, Y., and He, Q.Y. (2012). clusterProfiler: an R package for comparing biological themes among gene clusters. OMICS 16, 284–287.

Zhu, C., Yu, M., Huang, H., Juric, I., Abnousi, A., Hu, R., Lucero, J., Behrens, M.M., Hu, M., and Ren, B. (2019). An ultra high-throughput method for single-cell joint analysis of open chromatin and transcriptome. Nat Struct Mol Biol 26, 1063–1070.

Ziffra, R.S., Kim, C.N., Ross, J.M., Wilfert, A., Turner, T.N., Haeussler, M., Casella, A.M., Przytycki, P.F., Keough, K.C., Shin, D., et al. (2021). Single-cell epigenomics reveals mechanisms of human cortical development. Nature 598, 205–213.

